# BenchRep-T: A Systematic Evaluation of T-Cell Repertoire-Based Disease Diagnostics

**DOI:** 10.64898/2026.06.09.727013

**Authors:** Alejandro Buendia, Chiho Im, Anshul Kundaje, Scott D. Boyd, Liel Cohen-Lavi

## Abstract

Adaptive immune receptor repertoire sequencing has emerged as a promising modality for disease diagnosis, relying on computational methods to analyze the T-cell receptor (TCR) sequences in an individual’s blood sample. Published methods rely on different cohorts, preprocessing pipelines, and evaluation metrics, making direct comparison challenging. We present BenchRep-T, a benchmark that harmonizes 2,352 public TCR*β* repertoires from ten cohorts and eight conditions with fixed data splits, and evaluates nine methods spanning statistical enrichment of shared sequences, feature-engineered ensembles, deep multiple instance learning (MIL), and sequence clustering. We assess methods on four tasks: disease classification within and across cohorts, performance under reduced sequencing depth, recovery of known antigen-specific driver sequences, and sensitivity to demographic confounding. No single method dominated across all tasks. Feature-engineered ensembles, led by gradient-boosted trees on V/J-gene usage and short sequence motifs, matched or exceeded deep MIL methods when evaluated within cohorts, and were the most robust to reduced sequencing depth among the methods tested. However, high within-cohort performance did not imply transfer across cohorts, recovery of known antigen-specific TCRs, or robustness to demographic confounding. We release the harmonized data, splits, and code to support rigorous and reproducible evaluation of TCR repertoire classification methods and accelerate the development of immune repertoire-based diagnostics.

## 1 Introduction

T and B lymphocytes are the principal cells of the adaptive immune system, with each cell expressing a unique surface receptor—the T-cell receptor (TCR) or B-cell receptor (BCR)—generated by recombination of gene segments. Upon binding an antigen, such as a fragment of a pathogen protein, the cell proliferates to generate a clone of daughter cells sharing identical or nearly identical genetically rearranged receptors, together referred to as a clonotype. The adaptive immune receptor repertoire (AIRR) thus encodes a record of past and ongoing immune exposures, providing biomarkers for diverse clinical states [1–3].

Immune receptor repertoires are highly diverse both within and between individuals. The potential sequence space exceeds 10^18^, with each individual sampling approximately 10^8^–10^10^ distinct receptors of a given type. Overlap between individuals is limited: only a small fraction of receptors are “public” sequences shared across individuals, even among people with similar immunological exposures [1, 2]. Repertoire composition is also shaped by host genetics and by demographic factors including age and sex [1, 2, 4–7]. AIRR data has been challenging to generate at scale, requiring experimental and computational infrastructure, and most studies have modest cohort sizes relative to the underlying complexity of the repertoire. High-throughput sequencing of the TCR *β* chain (TCR*β*) has nonetheless identified disease-associated signatures across chronic viral infection [7], autoimmunity [8, 9], tuberculosis progression [10], and acute viral infection or vaccination [11, 12], suggesting that TCR*β* repertoires can support disease-specific prediction tasks.

Machine learning on AIRR data addresses two major prediction settings [2, 13]. In *receptor-level* prediction, models take an individual receptor, optionally together with a candidate epitope, and predict receptor–epitope binding. For TCRs, specificity and binding prediction have been assessed in community challenges and systematic comparisons of dozens of published models [14–18]. Here, we address *repertoire-level* prediction, where the input is a set of receptors from a biological sample, and the target is a sample-level label, such as disease status. This is a multiple instance learning problem, where each repertoire is a bag of sequences labeled only at the bag level [19–21]; the signal is carried by a small, typically unknown subset of sequences having limited overlap between individuals [7, 13, 22, 23].

AIRR classification often uses receptor V- and J-gene segment composition and complementarity-determining region 3 (CDR3) sequences. CDR3 is the most sequence-diverse region of each receptor, generated by gene-segment recombination and nucleotide insertion and deletion at the junctions, and is a major determinant of the recognition of a specific target antigen [1, 2]. AIRR datasets are also affected by the experimental protocols chosen for library generation, as well as biological variation [13]. Repertoire classifiers fall into four broad families (Figure 1): statistical and motif-based methods that identify disease-associated clonotypes or sequence motifs [7, 24, 25], deep multiple instance learning models [19, 26, 27], clustering methods that score repertoires by membership in disease-associated sequence clusters [28, 29], and ensembles that combine sub-models built on several complementary representations [12].

**Figure 1:**
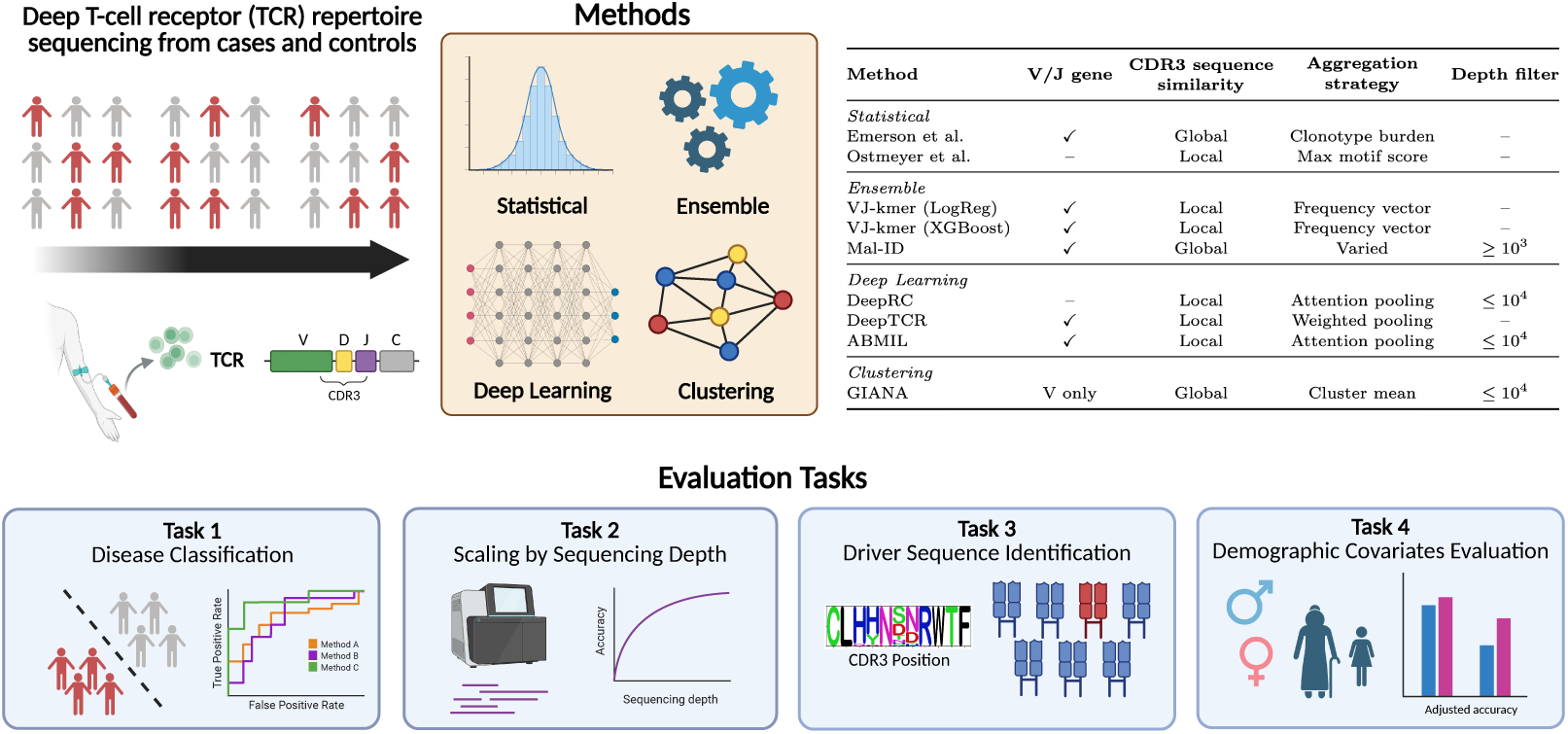
BenchRep-T overview. TCR*β* repertoires from cases and controls of eight disease states in ten cohorts are used to evaluate methods from four families on four tasks (**bottom**). The table (**top right**) lists each method’s use of V/J genes, CDR3 comparison (*global*: whole-sequence identity, distance, protein language model embedding, or alignment; *local*: sub-sequence features such as biochemical motifs, gapped *k*-mers, or convolutional windows), repertoire-level aggregation, and training-time depth filter (if any). Schematic created with BioRender.com.

Prior comparisons of TCR repertoire classifiers have relied largely on simulated repertoires with implanted signals and on the cytomegalovirus (CMV) cohort of Emerson et al. [7], varying properties such as signal frequency and training set size [19, 20, 25, 30]. These reached mixed conclusions: *k*-mer–based models were robust and competitive in some comparisons [20, 25], whereas deep attention-based models performed best in others [19]. The immuneML framework [30, 31] supports specifying, running, and sharing such analyses and implements three of the methods benchmarked here, and the AIRR-ML-25 challenge [32] scored submitted models on repertoire classification and label-associated sequence ranking across real and simulated datasets.

Extending these comparisons across more diseases, cohorts, sequencing platforms, and method families is important to clarify how model performance generalizes. Frameworks such as immuneML make such comparisons possible but, by design, leave the data, preprocessing, and splits to each user; a benchmark fixes them so that results are comparable across methods and studies. To this end, we present **BenchRep-T**, a benchmark for TCR*β*-based disease diagnostics (Figure 1). BenchRep-T harmonizes 2,352 public TCR*β* repertoires from 2,264 individuals across ten cohorts and eight conditions: five disease and vaccine cohorts with shared healthy controls from Zaslavsky et al. [12], and five immunoSEQ cohorts covering type 1 diabetes (T1D) [8, 9], tuberculosis progression [10], rheumatoid arthritis [33], and CMV serostatus [7]. We evaluate nine published or custom-designed TCR repertoire classification methods under identical inputs and preassigned data splits, on four tasks: (i) *disease classification*, within and across cohorts, (ii) *sequencing-depth scaling*, after downsampling repertoires to fixed depth, (iii) *driver sequence identification*, ranking known antigen-specific TCRs [11, 34], and (iv) *demographic-confounding analysis*, testing whether performance reflects demographic structure rather than repertoire-derived disease signal. We also compare runtime of included methods and their CPU and GPU memory use.

In summary, our main contributions are: (i) the benchmark itself, with harmonized data, fixed splits, and code for all methods, also released in an immuneML-compatible format; (ii) an evaluation protocol with bootstrap uncertainty estimates, paired significance tests, and demographics-only baselines; and (iii) a systematic comparison of nine methods across the benchmark tasks, which shows that no single method dominates and that high within-cohort performance does not imply transfer across cohorts, recovery of known antigen-specific TCRs, or robustness to demographic confounding.

## 2 Datasets and Preprocessing

BenchRep-T uses TCR*β* repertoire data from multiple sources [12, 35], all derived from peripheral blood. Repertoires are converted to a single AIRR schema (CDR3 amino-acid sequence, V gene, J gene) with matched gene nomenclature, CDR3 boundaries, and quality filters, so that every method receives the same input files (Appendix C.1). Age, sex, and self-reported ancestry are available for a subset of the cohorts (Figure 5a), and an HLA-DR risk-genotype category for Rawat T1D.

### 2.1 TCR Repertoire Data Sources

#### Mal-ID cohorts

The Zaslavsky et al. Mal-ID [12] study aggregates bulk BCR (IgH) and TCR*β* immunosequencing data from multiple clinical studies and medical centers, generated with a standardized sequencing protocol: TCR*β* was PCR-amplified from cDNA with gene-specific primers and sequenced on Illumina MiSeq. We use its 550 TCR*β* specimens from 542 participants, comprising 71,302,390 productive rearrangements after preprocessing, from 197 healthy/background controls, 98 **HIV**, 96 **T1D**, 64 **Lupus**, 58 **COVID-19**, and 37 **Influenza vaccine** specimens.

#### immunoSEQ cohorts

We include five publicly available TCR*β* datasets generated from genomic DNA with the Adaptive immunoSEQ assay [35], which differs from the Mal-ID protocol in PCR primers and initial data processing. A total of 216,302,987 productive rearrangements after preprocessing are present in these data, with one specimen per participant unless noted: (i) **T1D** Mitchell et al. [9], 196 specimens (171 T1D, 25 healthy controls); (ii) **T1D** Rawat et al. [8], 614 specimens from cohort 1 of the study (426 T1D, 188 unrelated, islet-autoantibody-negative controls; 684 relatives and 95 autoantibody-positive non-diabetic participants were excluded); (iii) **Tuberculosis (TB) progression** [10], 140 longitudinal specimens from 67 latently infected individuals (63 from 27 progressors, 77 from 40 non-progressing controllers); (iv) **Rheumatoid Arthritis (RA)** [33], 91 specimens from 84 individuals after excluding three low-depth repertoires (71 from 64 RA patients, 20 from 20 healthy controls); and (v) **CMV serostatus** Emerson et al. [7], 641 specimens from a discovery cohort (289 CMV^+^, 352 CMV*^−^*) and 120 specimens from a validation cohort (51 CMV^+^, 69 CMV*^−^*).

### 2.2 Disease Coverage and Evaluation Protocol

T1D appears in three cohorts. We combined the Mal-ID and Mitchell T1D specimens, which share no individuals (Appendix C.12), into a single dataset, so that each method is trained and tested on a mix of both assays and experimental batches. We evaluate Rawat T1D separately, and also test transfer between the T1D cohorts (Section 4.1). All datasets except CMV are partitioned into three participant-disjoint cross-validation folds, reused for every task: the three-fold design is inherited from the Mal-ID release, whose folds we reuse unchanged [12], and the Mitchell T1D, Rawat T1D, TB, and RA cohorts are split into three label-stratified folds to match. Rawat T1D is evaluated on its held-out fold 2, with model selection by nested cross-validation on folds 0 and 1. CMV keeps the original design [7], training on the discovery cohort and testing on the validation cohort.

Each model is trained as a binary classifier of cases against the cohort’s comparator group: healthy/background controls (Mal-ID diseases, pooled T1D, RA), unrelated islet-autoantibody-negative controls (Rawat T1D), CMV-seronegative donors (CMV), and non-progressing controllers (TB progressors). We report area under the receiver operating characteristic curve (AUROC) and area under the precision–recall curve (AUPRC) on pooled out-of-fold predictions, or on the held-out set for Rawat T1D and CMV.

## 3 Benchmarked Methods

We benchmark nine methods from the four families in Figure 1. Emerson et al., Ostmeyer et al., and Mal-ID are reimplemented from their published descriptions; DeepRC, DeepTCR, and GIANA use the authors’ released code with adapted data loading; and ABMIL and both VJ-kmer variants are new to this work. All methods run in a common codebase on the same inputs and splits (Section 2). Full descriptions and reproduction checks are provided in Appendices C.2, C.3, and C.4.

### Statistical and motif-based. Emerson et al

[7] selects (V gene, J gene, CDR3) clonotypes enriched in cases by a one-sided Fisher’s exact test, and scores each specimen with class-conditional beta-binomial models of its count of these clonotypes, conditioned on repertoire size. **Ostmeyer et al.** [24] encodes each CDR3 4-mer by its Atchley factors and relative abundance, and trains a logistic regression in which each repertoire is scored by its top-scoring 4-mer.

**Ensembles. VJ-kmer (Reg)** combines two *L*_1_-penalized logistic regression models, one on V/J-gene usage frequencies and one on CDR3 4-mer and gapped 4-mer frequencies, using a tuned weighted average; **VJ-kmer (XG)** replaces each base learner with a gradient-boosted tree classifier (XGBoost) [36]. **Mal-ID** [12] combines three sub-models based on complementary repertoire representations: (i) a model based on V/J gene usage frequencies, (ii) a convergent-clustering model that clusters sequences sharing V gene, J gene, and CDR3 length by CDR3 similarity and trains a classifier on counts of disease-enriched clusters, and (iii) a protein language model pipeline that embeds CDR3s with ESM-2 [37] and aggregates V-gene-specialized sequence-level scores to the specimen level.

### Deep multiple instance learning

As a learned counterpart to VJ-kmer, we implement an **Attention-Based Multiple Instance Learning (ABMIL) model** [27] that replaces hand-designed features with end-to-end learned embeddings. It encodes each CDR3 with a CNN combined with learned V/J-gene embeddings and pools these per-sequence features by gated attention. **DeepRC** [19] uses CDR3 sequences only: one-hot amino-acid and positional features are encoded with a one-dimensional CNN, and the sequence embeddings are pooled by an attention-weighted average, inspired by modern Hopfield networks, that learns to upweight disease-associated sequences. **DeepTCR** [26] combines a convolutional CDR3 encoder with learned V/J-gene embeddings, maps each unique sequence into a learned concept space, and represents the repertoire as the average of its per-sequence concept activations.

**Clustering. GIANA** [28] clusters TCRs by CDR3 and V gene using a BLOSUM-derived isometric embedding. Clusters are built from training sequences only; each is assigned the fraction of its training sequences that come from cases. Test sequences are assigned to these clusters by nearest-neighbor search, and a test repertoire is scored by the mean case fraction over its assigned sequences.

## 4 Evaluation Approaches and Results

### 4.1 Disease Classification Benchmark

We evaluate disease classification in nine within-cohort and two cross-cohort settings, each defined by the data a model is trained on and the data it is tested on. In the *within-cohort* settings, models are trained and tested on different individuals from the same source studies: seven single-cohort settings (HIV, Lupus, COVID-19, and Influenza from Mal-ID, plus RA, TB, and CMV) and two T1D settings (pooled Mal-ID and Mitchell, and held-out Rawat). In the *cross-cohort* settings, models trained on Rawat are applied without retraining to the Mal-ID and Mitchell T1D cohorts; T1D is the only condition profiled in three independent cohorts.

### Within-cohort performance

Performance varied substantially across methods and cohorts (Figure 2a; bootstrap standard errors in Tables B.1 and B.2; 3-fold cross-validation results for the two held-out cohorts, Rawat T1D and CMV, in Table B.3). The ensembles ranked at or near the top: VJ-kmer (XG) had the highest mean AUROC (0.81) and AUPRC (0.80) across the nine within-cohort settings, followed by Mal-ID (0.79 and 0.78), although it may benefit from a home-field advantage on the Mal-ID cohorts it was developed on. ABMIL and DeepTCR were competitive in AUROC in several settings, but their AUPRC was less consistent than that of the ensembles, and Ostmeyer et al. and GIANA had the lowest mean AUROC (0.61 for both). In paired comparisons (Table B.4; Appendix C.5), VJ-kmer (XG) and Mal-ID each significantly outperformed DeepRC in the five settings that include Mal-ID specimens (BH-adjusted *q <* 0.05, paired bootstrap and DeLong tests), whereas their differences from DeepTCR were setting-dependent. On Rawat T1D, both outperformed DeepTCR but not DeepRC, and no comparison was significant on RA, TB, or CMV.

**Figure 2:**
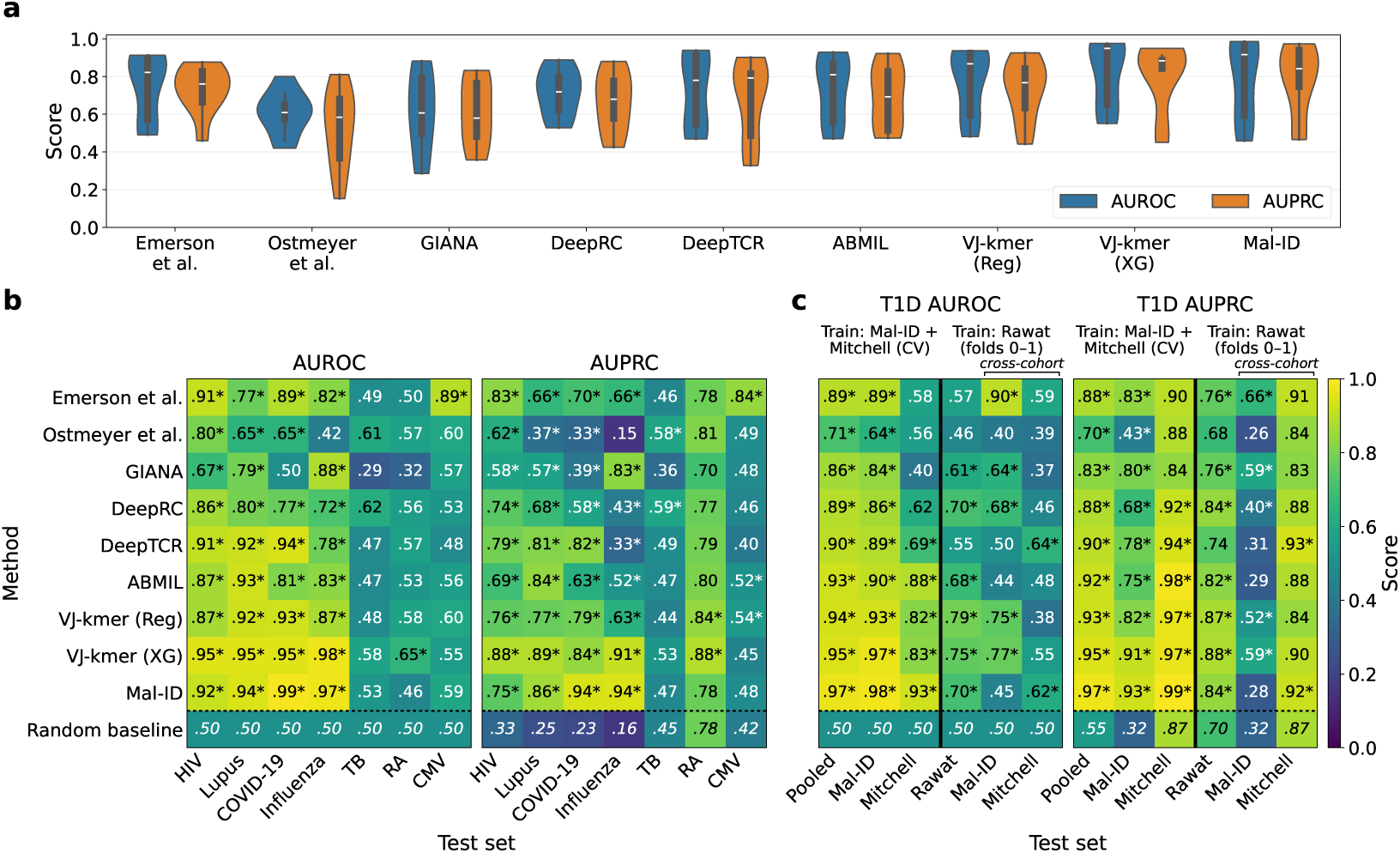
Disease classification performance. **(a)** AUROC (blue) and AUPRC (orange) of each method across the nine within-cohort settings; inner boxes mark the median and interquartile range. **(b)** AUROC and AUPRC for the seven single-cohort settings. **(c)** AUROC and AUPRC for the four T1D settings. In each heatmap, the first three columns (“Train: Mal-ID + Mitchell (CV)”) show the pooled within-cohort setting, trained under 3-fold cross-validation on Mal-ID and Mitchell and scored on all out-of-fold predictions (Pooled) and separately on each cohort’s specimens. The last three columns (“Train: Rawat (folds 0–1)”) show models trained on Rawat folds 0–1, scored on the held-out Rawat fold 2 (within-cohort) and, without retraining, on the full Mal-ID and Mitchell T1D cohorts (two cross-cohort settings). Within-cohort metrics use pooled 3-fold out-of-fold predictions, except Rawat T1D (fold 2) and CMV (trained on the Emerson discovery cohort, tested on its validation cohort). Bottom rows give the random baseline (0.5 for AUROC, the case fraction for AUPRC); asterisks mark estimates whose 95% participant-level bootstrap confidence interval lies above it.

Figure 2b breaks performance down for the seven single-cohort settings. By AUROC, VJ-kmer (XG) was best on HIV (0.953), Lupus (0.951), and RA (0.654) and tied with Mal-ID on Influenza (0.976 vs. 0.974; Mal-ID was best by AUPRC), and Mal-ID was best on COVID-19 (0.986). Only Emerson et al. performed well on CMV (AUROC 0.893 vs. at most 0.60 for all others), likely because the method was developed on this cohort [7]. RA and TB were the hardest conditions and do not separate the methods: in AUROC, only VJ-kmer (XG) exceeded chance on RA, and no method did on TB. GIANA’s below-chance AUROCs on RA and TB (0.325 and 0.286) likely reflect its fixed-direction cluster-enrichment score, which inverts when controls are more public or convergent than cases, and Emerson et al. assigned every RA specimen the fallback score of 0.5 because no clonotype passed its enrichment threshold. The source studies suggest that weak disease signal in the repertoires limits performance on these conditions [10, 33], and the small cohorts (91 RA and 140 TB specimens) widen the uncertainty of each estimate.

To check that the deep MIL methods were not disadvantaged by undertuning, we ran a matched-budget hyperparameter search (eight diverse configurations per method, three random seeds). Averaged across seeds, tuned ABMIL, DeepTCR, and DeepRC did not exceed the best benchmark method in any of the nine within-cohort settings (Appendix C.6; Tables B.6 and B.7). Replacing ABMIL’s CDR3 encoder with frozen pretrained representations from the protein language model ESM-2 [37], the TCR language model TCR-BERT [38], or the TCR variational autoencoder TCR-VALID [39] changed AUROC by *−*0.15 to +0.06 depending on the setting and encoder, without closing the gap to the top methods in most settings (Appendix C.7; Table B.9). Rawat T1D was a notable exception: ABMIL with TCR-BERT (AUROC 0.742 *±* 0.037) improved on baseline ABMIL by 0.06 and fell within the range of the feature-engineered ensembles (AUROC 0.70–0.79).

### T1D within cohorts and cross-cohort transfer

The three T1D cohorts span two assays: Mal-ID repertoires were sequenced from cDNA, and Mitchell and Rawat from genomic DNA with the immunoSEQ assay (Section 2). *Within cohorts*, every method exceeded chance in the pooled Mal-ID and Mitchell setting (Figure 2c, “Train: Mal-ID + Mitchell” columns; Mal-ID best, AUROC 0.969), but every method scored lower on Mitchell than on Mal-ID test specimens, and only five of nine exceeded chance on Mitchell alone. On held-out Rawat individuals (Rawat column), the best method reached AUROC 0.79 (VJ-kmer (Reg)). *Across cohorts* (Mal-ID and Mitchell columns under “Train: Rawat”; Table B.10), five of nine methods exceeded chance on Mal-ID, led by Emerson et al. (0.90, versus at most 0.77 for the others), whereas transfer to Mitchell largely failed despite its shared assay with Rawat (DeepTCR best, 0.64; five of nine below 0.5). Method rankings did not consistently carry over: the two transfer leaders, Emerson et al. and DeepTCR, were indistinguishable from chance on held-out Rawat individuals (0.57 and 0.55). No method exceeded chance on both cohorts; among the ensembles, both VJ-kmer variants did so on Mal-ID (0.75 and 0.77) and the Mal-ID method on Mitchell (0.62), so their within-cohort lead held only partly across cohorts.

### 4.2 Sequencing Depth Scaling and Computational Resources Benchmark

Sequencing depth varies widely across specimens and cohorts (Figure A.1) and can affect both signal detection and predictive performance. Published methods handle it differently, for example by subsampling or by normalizing counts to relative abundances [7, 12, 19, 24, 28]. To test its effects, we reran Lupus and HIV classification after downsampling every repertoire to a common depth *D ∈ {*1k, 5k, 10k, 25k, 50k, 75k*}* unique sequences, using five nested subsampling replicates in which each smaller subsample is contained in the larger ones, for Emerson et al., DeepRC, both VJ-kmer variants, and Mal-ID (Appendix C.8).

The ensembles were the least sensitive to depth (Figure 3a): from 75k down to 1k sequences (5k for Mal-ID), the AUROC of both VJ-kmer variants and Mal-ID dropped by less than 0.04. Emerson et al. depended most strongly on depth, falling to chance at 1k on Lupus and still improving at 75k on HIV, and DeepRC was intermediate, with a non-monotonic trend. AUPRC followed the same pattern (Figure A.2). Among these five methods, the ensembles were therefore both the most accurate and the most depth-robust. However, with DeepRC as the only deep MIL method tested, this comparison does not generalize to deep MIL models as a family.

**Figure 3:**
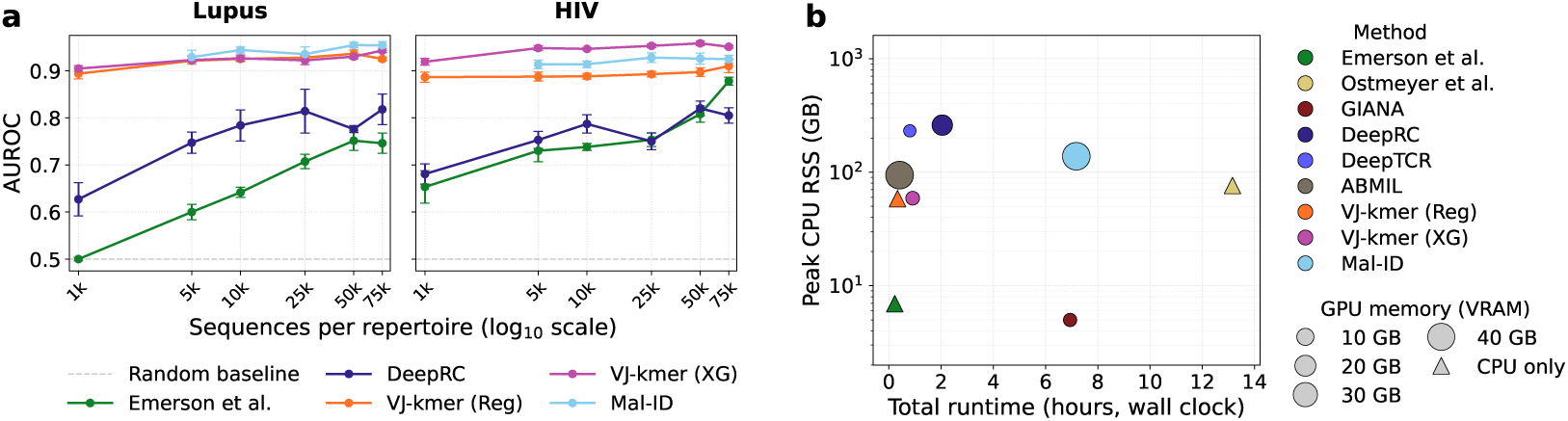
Sequencing depth scaling and computational requirements. **(a)** AUROC versus sequencing depth (unique sequences per repertoire) on Lupus (**left**) and HIV (**right**); markers show means and error bars show standard errors over five nested subsampling replicates. **(b)** End-to-end runtime (*x*-axis) and peak CPU memory (*y*-axis, log scale) for a single run on one HIV dataset fold; marker size indicates peak GPU memory, and triangles mark CPU-only methods.

We also measured end-to-end runtime and peak CPU and GPU memory for one HIV fold (Figure 3b; Appendix C.9). Emerson et al. and both VJ-kmer variants finished in under an hour, with only VJ-kmer (XG) using a GPU (3 GB), whereas GIANA and Mal-ID took about 7 h and Ostmeyer et al. 13 h. The deep MIL methods ran on GPUs in 0.4–2 h. Peak CPU memory ranged from 5–7 GB (GIANA, Emerson et al.) to 231–260 GB (DeepTCR, DeepRC), and ABMIL and Mal-ID used the most GPU memory (*∼*42 GB each).

### 4.3 Driver Sequence Identification

Accurate classification does not show that a method prioritizes biologically meaningful sequences. The *driver sequence identification* task therefore asks whether methods that produce per-sequence scores (Emerson et al., GIANA, DeepRC, ABMIL, and both VJ-kmer variants) rank known antigen-specific TCRs highly. Reference TCR*β* CDR3s with known specificity for HIV (1,747), COVID-19 (18,082), and Influenza (36,453) are taken from VDJdb [34] and, for COVID-19, also from Minervina et al. [11] (Appendix C.10). We label a specimen CDR3 as a *driver* if it has at least 90% Levenshtein similarity to a reference CDR3 for that disease. Within each fold, each method ranks the CDR3s of every held-out disease-positive specimen, and recall@*k* is the fraction of that specimen’s drivers among its top *k* sequences, macro-averaged over specimens (Appendix C.11). The reference sets are fixed, identical for all methods, and never used for training, and drivers are rare, making up 0.0004–0.03% (exact match) and 0.02–1.4% (fuzzy match) of each disease repertoire (Appendix C.12).

Emerson et al. most consistently ranks driver sequences highly across all three diseases (Figure 4). Its advantage is most pronounced at small and intermediate *k*: at *k*=1,000, Emerson et al. is clearly above the random baseline for all three diseases, whereas most other methods remain close to random or lower. GIANA performs relatively well for all diseases at *k*=1,000 and shows some enrichment at *k*=100 for HIV; however, it performs worse than random at *k*=10,000. At *k*=10,000 the gap between methods narrows, with Emerson et al., ABMIL, and VJ-kmer (Reg) recovering a slightly larger fraction of driver sequences than random for HIV and Influenza. For COVID-19, all methods perform close to random or worse. Absolute recall is highest for COVID-19 across all methods, but so is the random-chance baseline (e.g., 0.72 for COVID-19 vs. 0.32 for HIV and Influenza at *k*=10,000), so recall should be compared against each disease’s baseline rather than across diseases.

**Figure 4:**
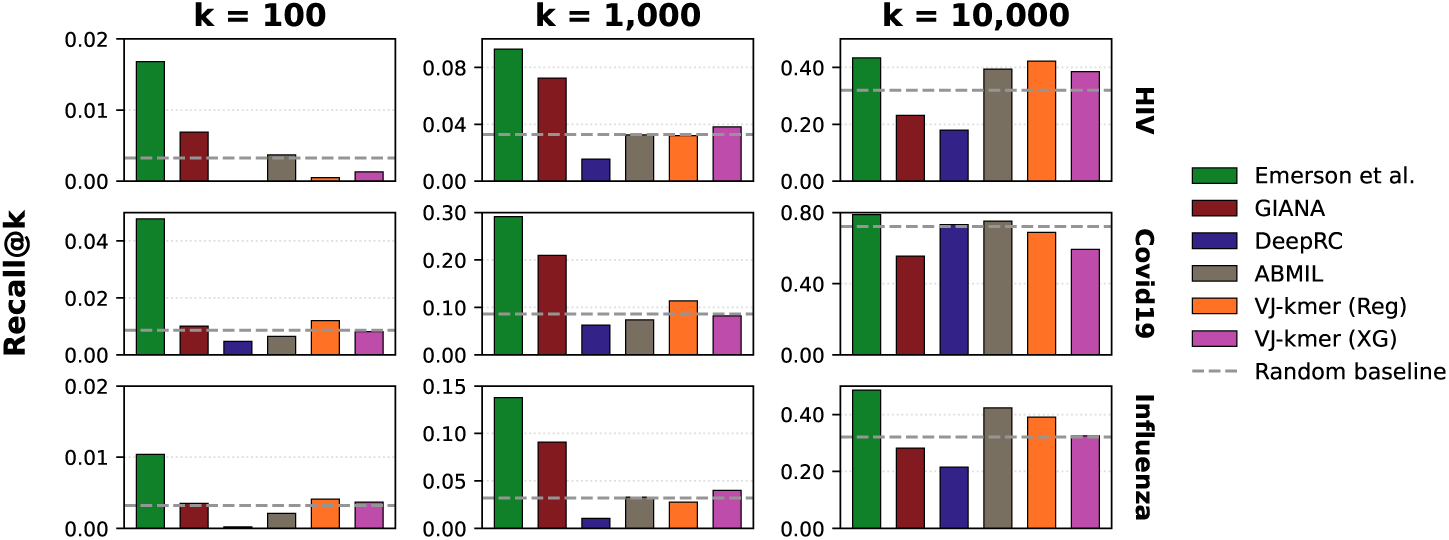
Macro-averaged specimen-level recall@*k* of driver sequences on HIV, COVID-19, and Influenza. Dashed lines show a random baseline: the mean recall@*k* of *k* sequences drawn uniformly at random from each disease-positive repertoire (10,000 draws), averaged across repertoires.

### 4.4 Demographic Confounding Analysis

TCR repertoire diversity and composition vary with age and genetic background, notably human leukocyte antigen (HLA) genes, which shape T-cell selection [5–7, 12, 24], and sex influences the risk of diseases such as lupus. Disease-associated and demographic signals may therefore be entangled through cohort selection or disease biology. For example, in the Mal-ID data, the Lupus cohort is predominantly female (81%), whereas 89% of the HIV cohort is of self-reported African ancestry (Figure 5a).

**Figure 5:**
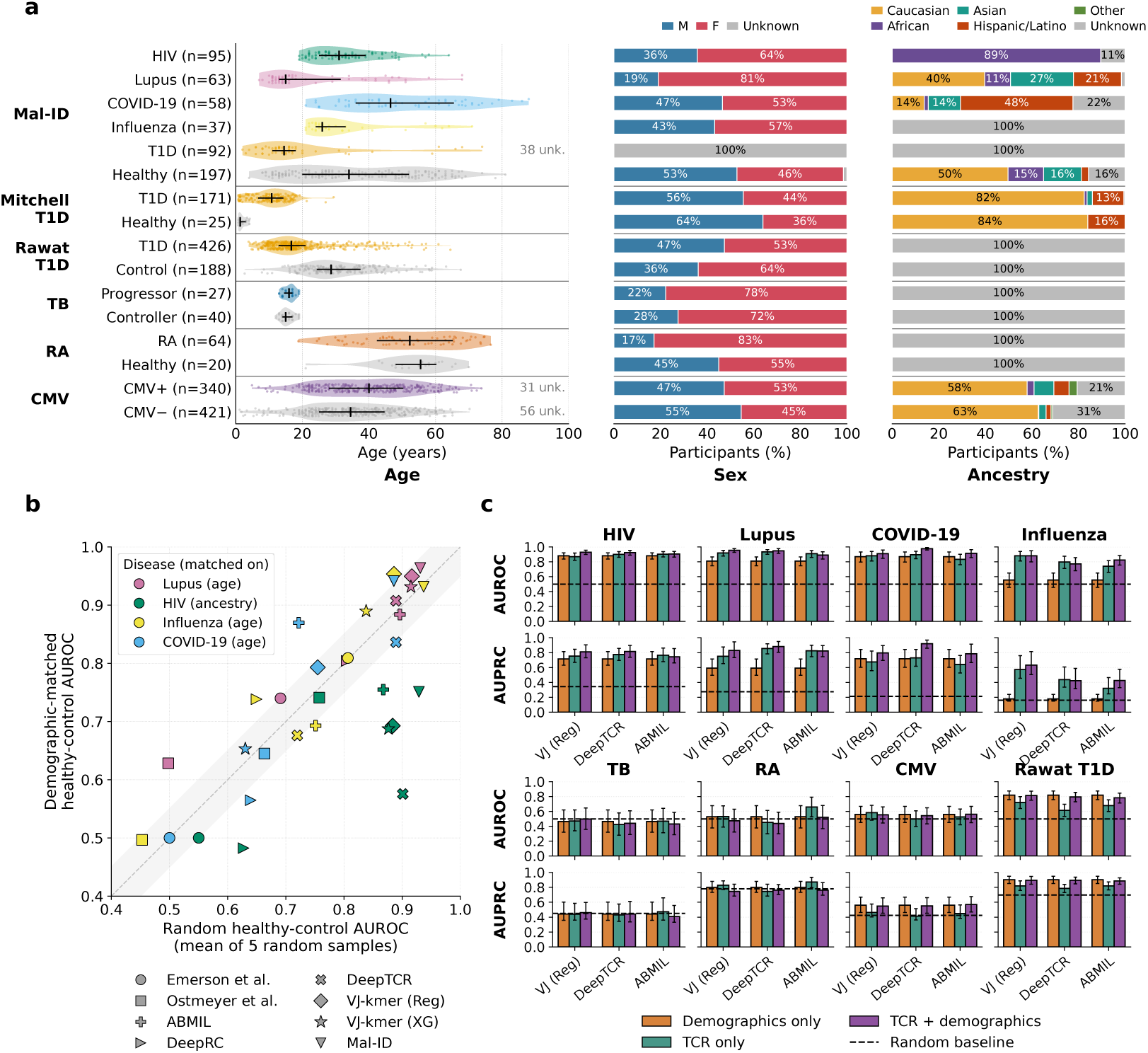
Demographic confounding. **(a)** Demographic composition of each cohort: age (violins), sex, and self-reported ancestry. Gray segments mark fields not recorded in the source metadata. **(b)** AUROC of eight methods with random healthy controls (*x*-axis, mean of five equally sized draws) versus controls matched on age or ancestry (*y*-axis). Points within the gray band differ by at most 0.05. **(c)** AUROC (top) and AUPRC (bottom) of a demographics-only logistic regression, three repertoire models (TCR only), and the same models with demographics appended (TCR + demographics), on specimens with complete demographic data. Rawat T1D uses held-out fold 2, CMV uses the validation cohort, and all others use pooled 3-fold out-of-fold predictions. Error bars are 95% participant-level bootstrap confidence intervals.

We probe how much of the apparent predictive signal reflects disease-associated repertoire features versus demographic imbalance with two complementary analyses (Appendix C.13). First, *demographic-matched disease classification* replaces the random pool of healthy controls with a subset matched to a dominant demographic confounder of the disease cohort: age for Lupus, Influenza, and COVID-19; for HIV, both cases and controls are restricted to specimens of self-reported African ancestry. Although the Lupus cohort is strongly sex-skewed, we match on age to vary only one demographic axis at a time, as the female skew reflects real-world lupus epidemiology. Each matched set is compared with an equally sized set of random healthy controls, averaged over five draws. Ancestry matching lowered HIV AUROC by 0.11–0.33 for six of eight methods (Figure 5b), consistent with the strong association between self-reported African ancestry and HIV status in this cohort. Age matching had mixed effects: on Lupus, AUROC was mostly unchanged or higher, whereas on Influenza and COVID-19 it moved in both directions.

Second, *demographic-aware disease classification* compares three repertoire models with a logistic regression on demographics alone and with the same models with demographics appended to their repertoire features. We evaluated VJ (Reg), the V/J-gene-usage sub-model of VJ-kmer (Reg), together with DeepTCR and ABMIL, in eight within-cohort settings, each restricted to specimens with complete demographic data (Figure 5c). Demographics alone were well above chance in Lupus, HIV, COVID-19, and Rawat T1D. In paired bootstrap tests of each repertoire-only model against the demographics-only baseline (24 tests, BH-corrected; Table B.11), all three repertoire models were significantly better on Lupus (demographics AUROC 0.81; ΔAUROC +0.10 to +0.13) and on Influenza, where demographics were near chance (AUROC 0.56; ΔAUROC +0.19 to +0.33). None was significantly better on HIV or COVID-19, and on Rawat T1D demographics alone (AUROC 0.82) significantly outperformed all three models (AUROC 0.61–0.72; ΔAUROC *−*0.10 to *−*0.21); age alone reproduces this baseline (AUROC 0.82; Table B.12), reflecting the age difference between cases and controls. On CMV, RA, and TB, demographics-only baselines were uninformative (AUROC 0.46–0.56), and no repertoire model differed significantly from them. Together, these results show that repertoire model performance should be interpreted relative to a demographics-only baseline, which can match or exceed repertoire models in some cohorts.

## 5 Discussion

By standardizing data, splits, and evaluation across methods, BenchRep-T reveals several broad patterns in how current methods extract disease signal from TCR repertoires. Within cohorts, no single method dominated across settings, underscoring the need to compare methods across multiple diseases and cohorts rather than draw conclusions from any single one. The VJ-kmer methods—ensembles based on V/J gene composition and gapped *k*-mer features—were nonetheless highly competitive with, and in several settings outperformed, methods that use deep representations. This suggests that much of the signal currently accessible in these cohorts lies in repertoire-level composition, which may be easier to learn than sequence-level patterns that deep multiple instance learning methods must learn from the small number of labeled repertoires per cohort. Replacing ABMIL’s encoder with pretrained embeddings and matched-budget hyperparameter tuning on the deep learning methods both gave cohort-dependent changes, but neither consistently closed the gap to the best ensemble method. Because the encoder ablation held ABMIL’s aggregation fixed, and the three encoders performed almost identically in four of the six settings where they exceeded chance (Table B.9), the remaining gap may lie partly in MIL aggregation rather than sequence representation alone.

Within-cohort rankings also did not carry over across cohorts. Rawat-trained T1D models transferred to Mal-ID for five of nine methods but largely failed on Mitchell despite its shared assay with Rawat. The methods that transferred best were near chance on held-out Rawat individuals, and no method exceeded chance on both cohorts. Because this evaluation covers a single training cohort and condition, it shows that rankings can change across cohorts rather than how often they do, but it cautions against extrapolating within-cohort rankings, including the ensembles’ lead, to new cohorts. Classification and driver recovery also ranked methods differently: Emerson et al. [7] recovered driver sequences better than several methods with higher AUROC. We hypothesize that this reflects the method’s focus on detecting identical sequences enriched among same-diagnosis participants, which resemble the driver reference sequences from the literature.

Depth and demographics shape these results in ways that classification scores alone do not reveal. Under downsampling, the ensembles were the least depth-sensitive of the five methods tested, likely because V/J gene usage is stable under subsampling and alone reaches high AUROC on the Mal-ID cohorts (Table B.13). Depth can differ between cases and controls: Mal-ID Influenza, COVID-19, and T1D repertoires are about fourfold shallower than the healthy controls (Figure A.1), so methods may partly exploit depth. Demographics alone matched the repertoire models on HIV and COVID-19 and outperformed them on Rawat T1D, but not on Lupus or Influenza, where the repertoire models carried signal beyond the measured demographics. On Rawat T1D, age alone reaches AUROC 0.82 (Section 4.4), so apparent transfer from Rawat-trained models may also partly reflect age rather than disease signal. Both within- and cross-cohort performance can therefore reflect cohort-specific confounders such as depth and age, and should be read against demographics-only baselines.

Several limitations remain, and they are also part of what the benchmark exposes about currently available TCR repertoire datasets. The data themselves are limited: cohorts are small, and several are demographically imbalanced (Figure 5a). The Mal-ID cohorts [12] contribute to five of the nine within-cohort settings and are the only source for four conditions (HIV, Lupus, Influenza, and COVID-19), and since the Mal-ID method was developed on these specimens, its performance on them may be optimistic. Two implementations differ slightly from their originals: Ostmeyer et al. uses 200 rather than 100,000–375,000 optimization restarts (which we found did not meaningfully affect performance), and our Emerson et al. reimplementation reaches AUROC 0.89 on the CMV validation cohort (95% CI 0.83–0.95), consistent with the published 0.94 (Appendix C.4). Finally, the evaluation is limited in scope. Cross-cohort models were applied to the target cohorts without correcting for technical differences between cohorts (e.g., sequencing platform or depth) or adapting the models to them, either of which might improve transfer. Moreover, we evaluate only binary classification, so strong performance here may not carry over to the multiclass diagnosis setting.

Many of these limitations reflect how repertoire cohorts are currently collected and shared, and overcoming them will require progress in data generation as well as in methods. We recommend that experimental groups (i) release data in a standardized, AIRR-compliant schema with fixed, published cross-validation splits, so that results are comparable across studies; (ii) include demographic and clinical metadata (age, sex, ancestry, and HLA) so that confounding can be measured and controlled; (iii) share both raw and processed, cleaned data with machine-readable metadata and a datasheet in persistent, open repositories rather than account-gated portals (when that is consistent with study participant informed consent); (iv) report sequencing depth and template counts, since methods differ in their robustness to depth and depth can differ between cases and controls; (v) expand and improve data quantity and quality by ensuring adequate sequencing depth and recruiting more independent participants, ideally across multiple centers, balancing on age, sex, and ancestry, so that cohorts pooled across studies become large enough to evaluate performance in relation to demographics and HLA genotype variation; (vi) record and release processing batch, sequencing run, and collection site; and (vii) where study design allows, process and sequence cases and controls in mixed batches, so that batch identity is not fully confounded with the disease label.

## Conclusion

We introduced BenchRep-T, a benchmark that harmonizes ten public TCR*β* repertoire cohorts and evaluates nine classification methods under identical preprocessing, splits, and uncertainty estimates. Within cohorts, feature-engineered ensembles were the strongest methods overall and the most depth-robust of the methods tested, with the VJ-kmer variants among the fastest to run. However, three findings caution against reading classification scores as evidence of generalizable disease signal: method rankings did not carry over under cross-cohort transfer, classification performance and recovery of known antigen-specific TCRs ranked methods differently, and demographics alone matched or exceeded repertoire models on several cohorts. These results suggest that progress in repertoire-based diagnostics currently depends as much on cohort design and data release as on model architecture. We release BenchRep-T as an extensible benchmark and set of datasets to encourage more rigorous and reproducible progress in immune repertoire-based disease diagnostics.

## Code and Dataset Availability

The code and documentation for our data processing and evaluations are available at https://github.com/boyd-lab/BenchRep-T. The Mal-ID re-implementation is available at https://github.com/liel-cohen/Mal-ID-Lite. The processed repertoire files for all experiments conducted in the paper are available at https://huggingface.co/datasets/lordim/BenchRep-T. The repository also hosts a Croissant metadata file with Responsible AI fields and a datasheet covering collection, consent, demographics, and licensing. BenchRep-T-authored metadata, data splits, and documentation are released under CC BY 4.0; the repertoire sequence data remain subject to the licenses and terms of use of their original sources (Mal-ID: CC BY-NC-SA 4.0; immuneACCESS cohorts: Adaptive immuneACCESS terms of use), which we credit and preserve in the release.

## Author Contributions

A.B. and C.I. initiated the project. A.B., C.I., and L.C.-L. designed the experiments with input from A.K. and S.D.B. A.B. adapted the GIANA, DeepRC, and DeepTCR models, and implemented the VJ-kmer regression, VJ-kmer XGBoost, and ABMIL methods. C.I. reimplemented the Emerson et al. and Ostmeyer et al. methods. L.C.-L. reimplemented the Mal-ID method. C.I., L.C.-L., and A.B. preprocessed the data used in the benchmarks, and conducted analysis and plotting of results. A.B. ran the encoder ablations, hyperparameter tuning experiments, and computational resources benchmark. C.I. and A.B. ran the statistical significance analyses. All authors wrote and revised the manuscript. L.C.-L., A.K., and S.D.B. supervised the project.

## Acknowledgments and Declaration of Interests

A.B. acknowledges support from the Warren Alpert Computational Biology & Artificial Intelligence Fellowship, the MAC3 Computing in Precision Health Fellowship, and NIH training grant T15LM007033. C.I. acknowledges support from the Stanford Bio-X Fellowship. A.K. is on the scientific advisory board of Immunera, SerImmune, and TensorBio; is a consultant with Bristol Myers Squibb and Inari; was a consultant with Illumina; and has a financial stake in Immunera, TensorBio, DeepGenomics, SerImmune, Immunai, and Freenome. S.B. declares grants from NIH (U19AI057229, U19AI67903, U54CA260517, U19AI090019, R01AI130398, R01AI127877), and the Bill and Melinda Gates Foundation; philanthropic gifts; consulting for Regeneron, Sanofi, Novartis, Genentech, Pfizer, Visterra, and Otsuka; stock ownership in AbCellera Biologics; and being a scientific cofounder of Immunera.

## A Supplementary Figures

**Figure A.1:**
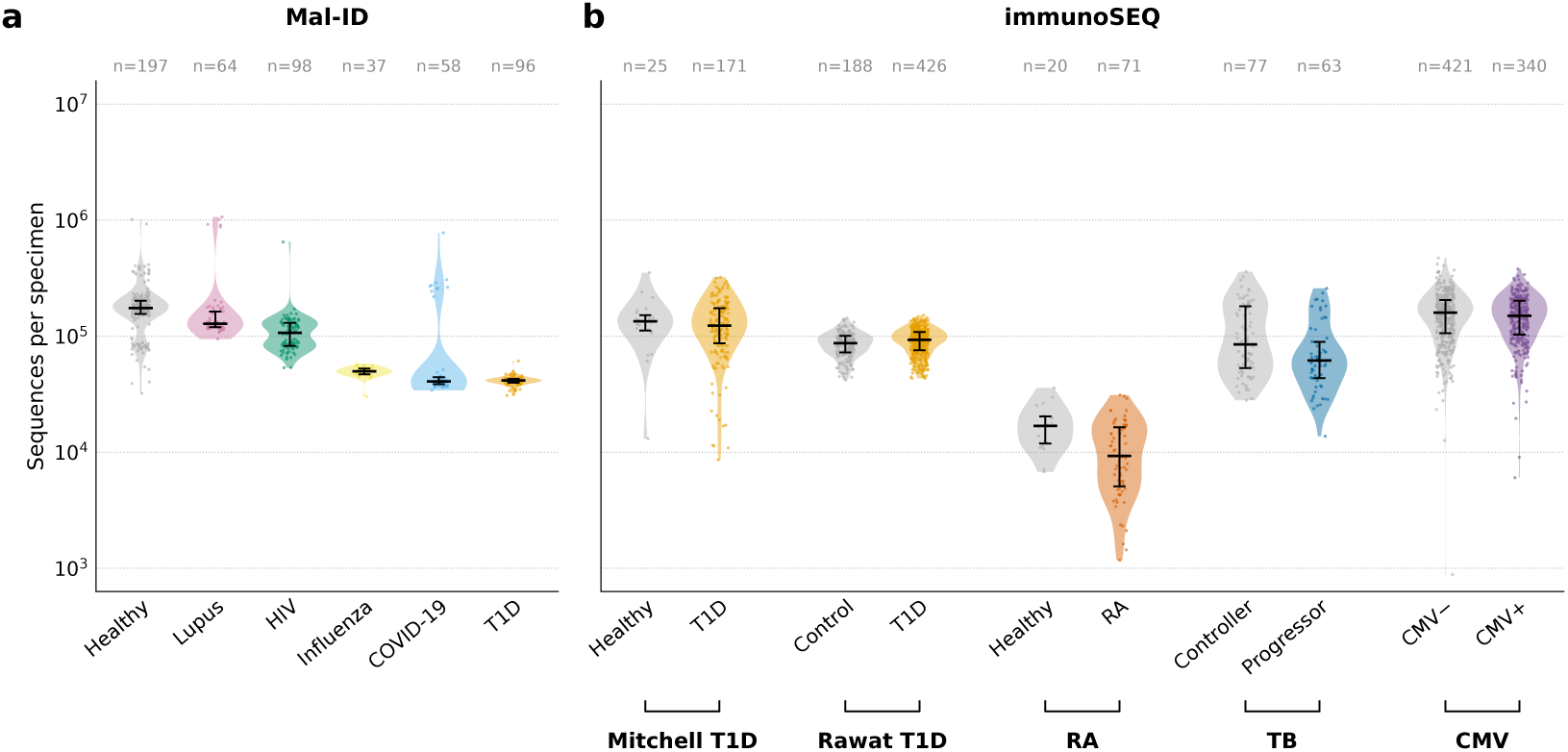
Repertoire sequencing depth distribution of the (a) Mal-ID and (b) immunoSEQ cohorts.

**Figure A.2:**
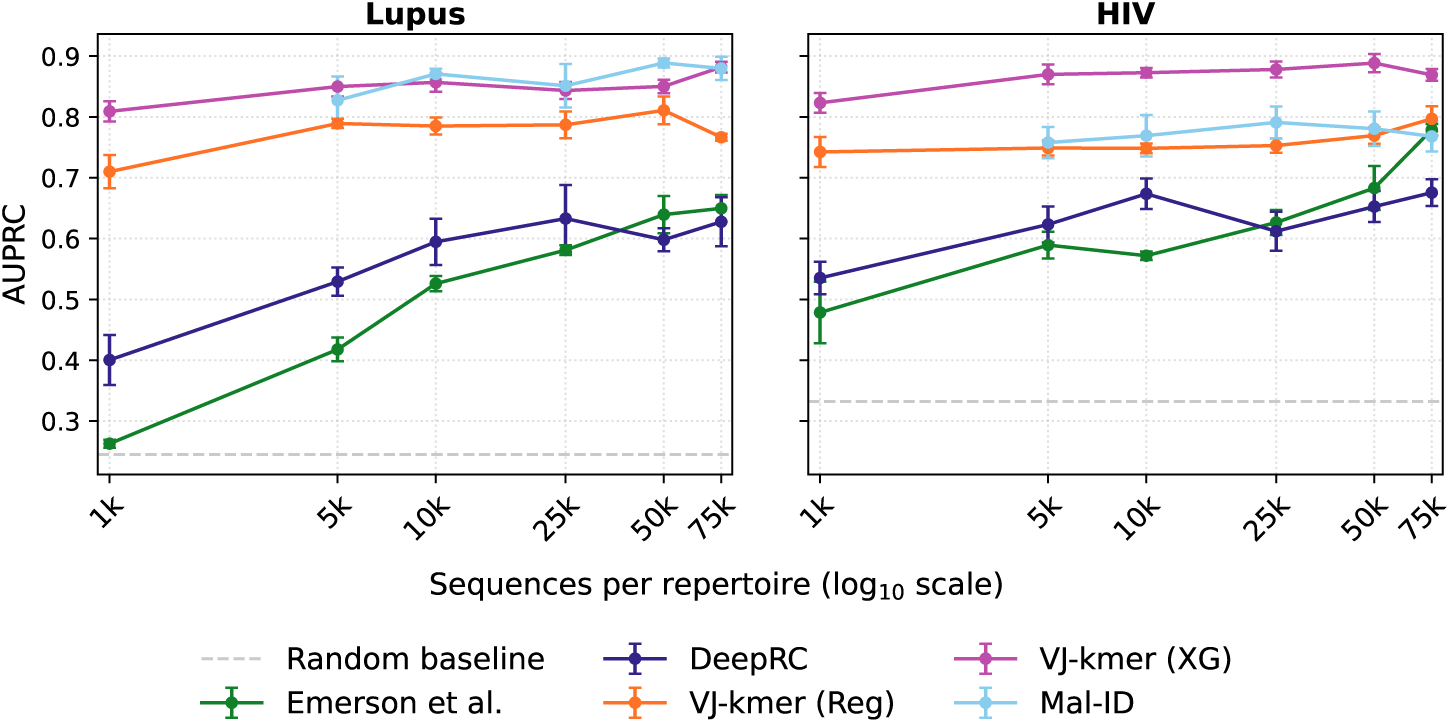
Disease-classification performance as a function of sequencing depth for Lupus (**left**) and HIV (**right**). Markers show mean AUPRC across five nested subsampling replicates, with error bars denoting the standard error across replicates. Mal-ID is evaluated from 5k sequences upward.

## B Supplementary Tables

**Table B.1:** Disease classification AUROC for all methods and settings. Point estimates with participant-level bootstrap standard errors in parentheses (*B* = 10,000; Appendix C.5); the best method per setting is in bold. Settings marked *‡* are evaluated on a single held-out set (the Rawat fold-2 evaluation fold and the Emerson validation cohort); all others use pooled 3-fold out-of-fold predictions. The RA cohort has 91 specimens, but Mal-ID’s minimum-repertoire-size filter (at least 1,000 sequences) excludes one small sample, so all methods are scored on the remaining 90 for a paired comparison. Emerson et al. on RA assigns every specimen the fallback score 0.5, so its standard error is not applicable.

| Method | HIV<br>(*n* = 295) | Lupus<br>(*n* = 261) | COVID-19<br>(*n* = 255) | Influenza<br>(*n* = 234) | T1D (pooled)<br>(*n* = 489) | TB<br>(*n* = 140) | RA<br>(*n* = 90) | CMV<sup>‡</sup><br>(*n* = 120) | Rawat T1D<sup>‡</sup><br>(*n* = 204) |
|---|---|---|---|---|---|---|---|---|---|
| Emerson et al. | 0.913 (.017) | 0.774 (.040) | 0.886 (.023) | 0.822 (.042) | 0.894 (.016) | 0.491 (.070) | 0.500 (n/a) | **0.893** (.032) | 0.572 (.043) |
| Ostmeyer et al. | 0.800 (.026) | 0.647 (.040) | 0.648 (.038) | 0.420 (.049) | 0.712 (.023) | 0.609 (.058) | 0.572 (.060) | 0.597 (.052) | 0.461 (.042) |
| GIANA | 0.672 (.039) | 0.791 (.036) | 0.499 (.051) | 0.881 (.045) | 0.861 (.019) | 0.286 (.064) | 0.325 (.066) | 0.570 (.054) | 0.607 (.042) |
| DeepRC | 0.863 (.024) | 0.796 (.037) | 0.767 (.034) | 0.718 (.052) | 0.887 (.015) | **0.622** (.062) | 0.561 (.085) | 0.528 (.053) | 0.704 (.039) |
| DeepTCR | 0.910 (.017) | 0.920 (.020) | 0.939 (.015) | 0.780 (.038) | 0.902 (.014) | 0.469 (.084) | 0.569 (.078) | 0.484 (.053) | 0.550 (.044) |
| ABMIL | 0.865 (.022) | 0.927 (.018) | 0.809 (.037) | 0.829 (.038) | 0.929 (.012) | 0.470 (.084) | 0.528 (.076) | 0.560 (.055) | 0.679 (.039) |
| VJ-kmer (Reg) | 0.867 (.023) | 0.923 (.019) | 0.928 (.017) | 0.872 (.038) | 0.937 (.012) | 0.481 (.072) | 0.575 (.070) | 0.597 (.052) | **0.791** (.037) |
| VJ-kmer (XG) | **0.953** (.011) | **0.951** (.014) | 0.950 (.015) | **0.976** (.009) | 0.949 (.010) | 0.579 (.075) | **0.654** (.064) | 0.551 (.054) | 0.755 (.035) |
| Mal-ID | 0.916 (.016) | 0.945 (.018) | **0.986** (.006) | 0.974 (.017) | **0.969** (.007) | 0.528 (.070) | 0.459 (.072) | 0.592 (.053) | 0.701 (.040) |

**Table B.2:**
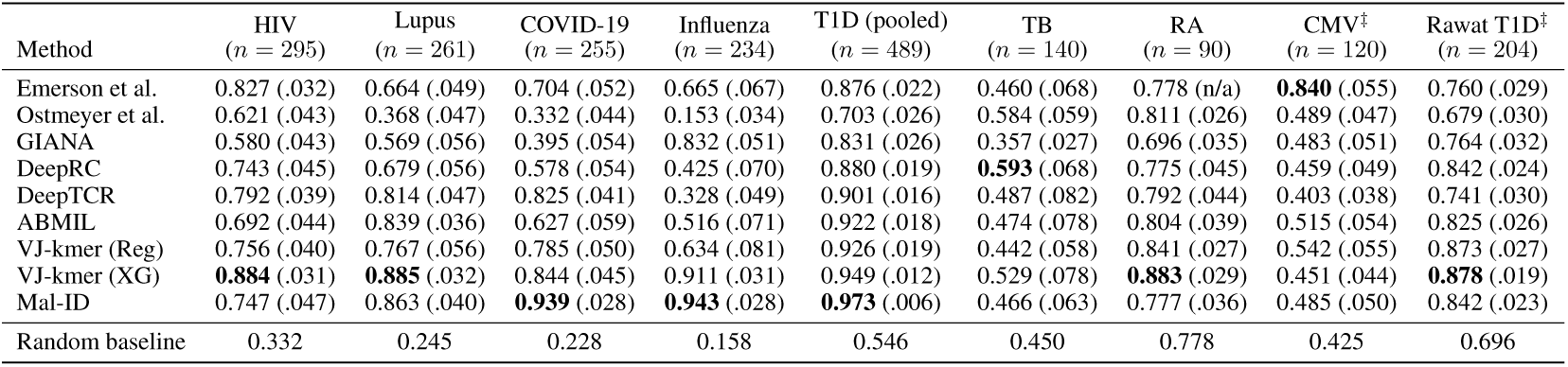
Disease classification AUPRC for all methods and settings, with the random (positive-class rate) baseline. Point estimates with participant-level bootstrap standard errors in parentheses (*B* = 10,000; Appendix C.5); the best method per setting is in bold. Settings marked *‡* are evaluated on a single held-out set (the Rawat fold-2 evaluation fold and the Emerson validation cohort); all others use pooled 3-fold out-of-fold predictions. Emerson et al. on RA assigns every specimen the fallback score 0.5, so its standard error is not applicable.

**Table B.3:** Three-fold cross-validation on the Emerson CMV and Rawat T1D cohorts. AUROC and AUPRC are computed on out-of-fold predictions pooled across the three folds, with participant-level bootstrap standard errors in parentheses (*B* = 10,000; Appendix C.5); the best method per column is in bold. For CMV, folds 0 and 1 partition the discovery cohort and fold 2 is the independent validation cohort; one discovery-cohort specimen with fewer than 1,000 sequences after preprocessing is excluded (Appendix C.1). The main-text results for these cohorts use the single held-out set (Tables B.1 and B.2).

|  | CMV (*n* = 760; 340 CMV<sup>+</sup>) |  | Rawat T1D (*n* = 614; 426 T1D) |  |
|---|---|---|---|---|
| Method | AUROC | AUPRC | AUROC | AUPRC |
| Emerson et al. | **0.768** (.017) | **0.745** (.019) | 0.550 (.025) | 0.735 (.018) |
| Ostmeyer et al. | 0.604 (.021) | 0.537 (.022) | 0.573 (.025) | 0.749 (.017) |
| GIANA | 0.508 (.021) | 0.452 (.018) | 0.585 (.025) | 0.744 (.019) |
| DeepRC | 0.532 (.021) | 0.477 (.020) | 0.635 (.024) | 0.802 (.016) |
| DeepTCR | 0.533 (.021) | 0.484 (.020) | 0.574 (.025) | 0.759 (.018) |
| ABMIL | 0.495 (.021) | 0.453 (.019) | 0.682 (.023) | 0.817 (.017) |
| VJ-kmer (Reg) | 0.528 (.021) | 0.497 (.021) | 0.759 (.021) | 0.862 (.015) |
| VJ-kmer (XG) | 0.529 (.021) | 0.468 (.018) | **0.783** (.020) | **0.870** (.016) |
| Mal-ID | 0.548 (.021) | 0.487 (.020) | 0.691 (.024) | 0.813 (.017) |
| Random baseline | 0.500 | 0.447 | 0.500 | 0.694 |

**Table B.4:** Paired comparisons of the feature-ensemble methods (VJ-kmer (XG), Mal-ID) against the deep-learning methods (DeepRC, DeepTCR) on all nine within-cohort settings (36 tests). ΔAUROC is AUROC(ensemble) *−* AUROC(deep) with its paired participant-bootstrap standard error (*B* = 10,000; Appendix C.5). *q*-values are Benjamini–Hochberg adjusted across the 36 tests, separately for the paired bootstrap and DeLong’s test; the two tests agree on all 36 calls at *q <* 0.05, with 18 significant. Settings marked *‡* are evaluated on a single held-out set (Rawat fold 2; Emerson validation cohort); all others use pooled 3-fold out-of-fold predictions.

| Setting | Comparison | AUROC<br>(ens. / deep) | ΔAUROC<br>± SE | *q*<br>(bootstrap) | *q*<br>(DeLong) |
|---|---|---|---|---|---|
| HIV<br>(*n* = 295) | VJ-kmer (XG) vs. DeepRC | 0.953 / 0.863 | +0.090 ± 0.023 | <0.001 | <0.001 |
|  | VJ-kmer (XG) vs. DeepTCR | 0.953 / 0.910 | +0.043 ± 0.015 | 0.007 | 0.012 |
|  | Mal-ID vs. DeepRC | 0.916 / 0.863 | +0.053 ± 0.023 | 0.032 | 0.044 |
|  | Mal-ID vs. DeepTCR | 0.916 / 0.910 | +0.006 ± 0.014 | 0.713 | 0.705 |
| Lupus<br>(*n* = 261) | VJ-kmer (XG) vs. DeepRC | 0.951 / 0.796 | +0.155 ± 0.034 | <0.001 | <0.001 |
|  | VJ-kmer (XG) vs. DeepTCR | 0.951 / 0.920 | +0.031 ± 0.017 | 0.110 | 0.120 |
|  | Mal-ID vs. DeepRC | 0.945 / 0.796 | +0.149 ± 0.038 | <0.001 | <0.001 |
|  | Mal-ID vs. DeepTCR | 0.945 / 0.920 | +0.025 ± 0.019 | 0.313 | 0.283 |
| COVID-19<br>(*n* = 255) | VJ-kmer (XG) vs. DeepRC | 0.950 / 0.767 | +0.184 ± 0.037 | <0.001 | <0.001 |
|  | VJ-kmer (XG) vs. DeepTCR | 0.950 / 0.939 | +0.011 ± 0.018 | 0.611 | 0.596 |
|  | Mal-ID vs. DeepRC | 0.986 / 0.767 | +0.219 ± 0.035 | <0.001 | <0.001 |
|  | Mal-ID vs. DeepTCR | 0.986 / 0.939 | +0.046 ± 0.015 | 0.002 | 0.004 |
| Influenza<br>(*n* = 234) | VJ-kmer (XG) vs. DeepRC | 0.976 / 0.718 | +0.257 ± 0.053 | <0.001 | <0.001 |
|  | VJ-kmer (XG) vs. DeepTCR | 0.976 / 0.780 | +0.196 ± 0.037 | <0.001 | <0.001 |
|  | Mal-ID vs. DeepRC | 0.974 / 0.718 | +0.256 ± 0.049 | <0.001 | <0.001 |
|  | Mal-ID vs. DeepTCR | 0.974 / 0.780 | +0.195 ± 0.033 | <0.001 | <0.001 |
| T1D (pooled)<br>(*n* = 489) | VJ-kmer (XG) vs. DeepRC | 0.949 / 0.887 | +0.061 ± 0.014 | <0.001 | <0.001 |
|  | VJ-kmer (XG) vs. DeepTCR | 0.949 / 0.902 | +0.047 ± 0.014 | 0.003 | 0.002 |
|  | Mal-ID vs. DeepRC | 0.969 / 0.887 | +0.082 ± 0.013 | <0.001 | <0.001 |
|  | Mal-ID vs. DeepTCR | 0.969 / 0.902 | +0.067 ± 0.011 | <0.001 | <0.001 |
| TB<br>(*n* = 140) | VJ-kmer (XG) vs. DeepRC | 0.579 / 0.622 | −0.044 ± 0.081 | 0.658 | 0.541 |
|  | VJ-kmer (XG) vs. DeepTCR | 0.579 / 0.469 | +0.109 ± 0.097 | 0.382 | 0.125 |
|  | Mal-ID vs. DeepRC | 0.528 / 0.622 | −0.094 ± 0.081 | 0.382 | 0.214 |
|  | Mal-ID vs. DeepTCR | 0.528 / 0.469 | +0.059 ± 0.097 | 0.611 | 0.440 |
| RA<br>(*n* = 90) | VJ-kmer (XG) vs. DeepRC | 0.654 / 0.561 | +0.094 ± 0.099 | 0.446 | 0.440 |
|  | VJ-kmer (XG) vs. DeepTCR | 0.654 / 0.569 | +0.086 ± 0.085 | 0.436 | 0.401 |
|  | Mal-ID vs. DeepRC | 0.459 / 0.561 | −0.102 ± 0.131 | 0.534 | 0.521 |
|  | Mal-ID vs. DeepTCR | 0.459 / 0.569 | −0.110 ± 0.099 | 0.382 | 0.385 |
| CMV<sup>‡</sup><br>(*n* = 120) | VJ-kmer (XG) vs. DeepRC | 0.551 / 0.528 | +0.023 ± 0.076 | 0.779 | 0.785 |
|  | VJ-kmer (XG) vs. DeepTCR | 0.551 / 0.484 | +0.067 ± 0.070 | 0.446 | 0.440 |
|  | Mal-ID vs. DeepRC | 0.592 / 0.528 | +0.064 ± 0.080 | 0.527 | 0.503 |
|  | Mal-ID vs. DeepTCR | 0.592 / 0.484 | +0.108 ± 0.063 | 0.159 | 0.143 |
| Rawat T1D<sup>‡</sup><br>(*n* = 204) | VJ-kmer (XG) vs. DeepRC | 0.755 / 0.704 | +0.050 ± 0.035 | 0.245 | 0.218 |
|  | VJ-kmer (XG) vs. DeepTCR | 0.755 / 0.550 | +0.205 ± 0.047 | <0.001 | <0.001 |
|  | Mal-ID vs. DeepRC | 0.701 / 0.704 | −0.003 ± 0.046 | 0.944 | 0.946 |
|  | Mal-ID vs. DeepTCR | 0.701 / 0.550 | +0.152 ± 0.055 | 0.022 | 0.012 |

**Table B.5:**
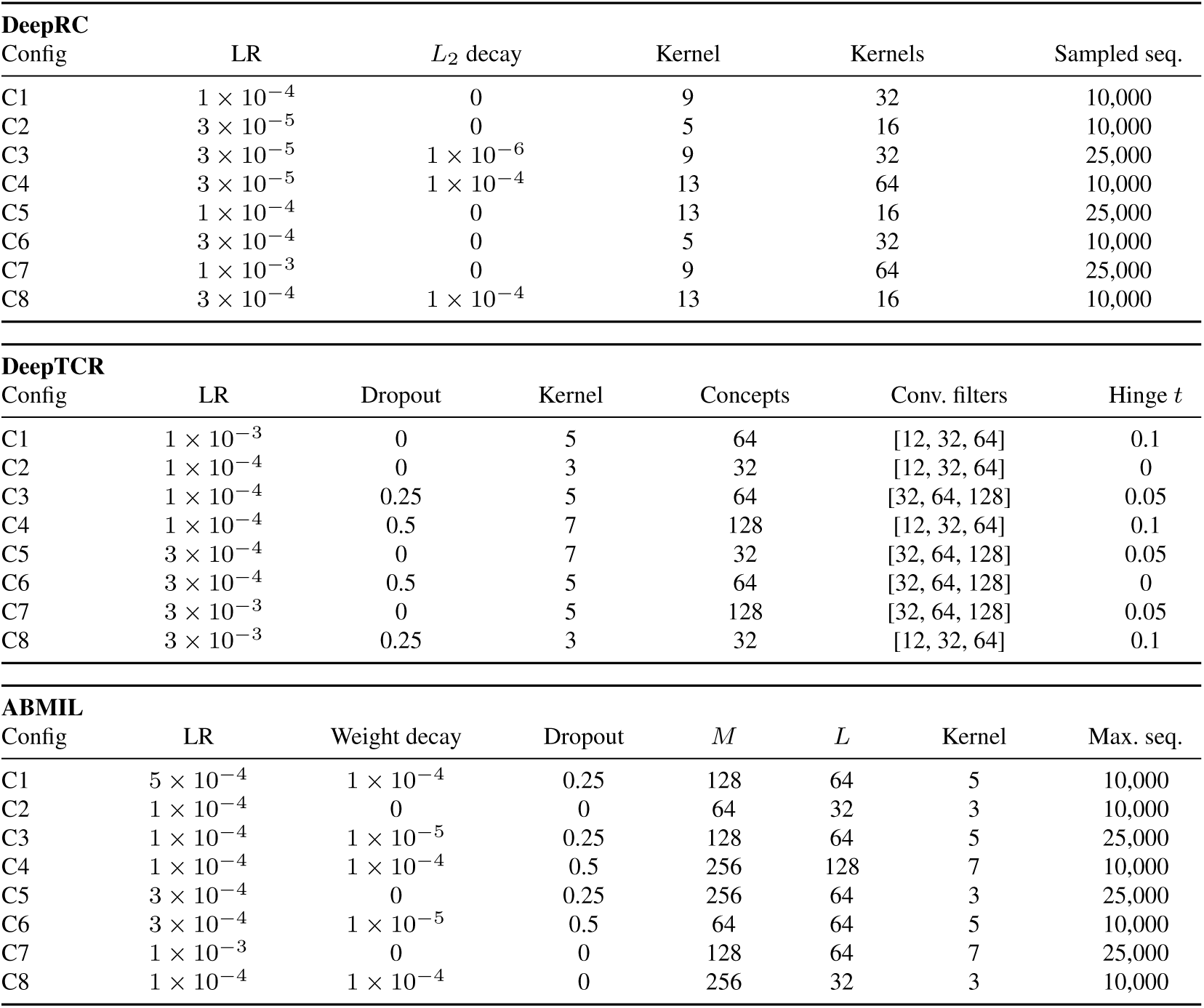
The eight hyperparameter configurations evaluated for each deep learning method in the matched-budget tuning (Appendix C.6). C1 is each method’s benchmark default configuration (Appendix C.2). DeepTCR convolutional filters are the layer widths of its small and medium networks.

**Table B.6:** Stage 1 of the matched-budget tuning: mean validation AUROC of each of the eight configurations (C1–C8; Table B.5) at seed 7, averaged over the tuning contexts (Appendix C.6). For the seven 3-fold settings the three contexts each train on one benchmark fold and validate on another; for settings marked *‡* the two contexts use folds 0 and 1 only, so the evaluation fold is never seen. The selected (highest-ranked) configuration for each method and setting is in bold; it is the configuration retrained and evaluated in Table B.7. C1 is each method’s benchmark default.

| Setting | Method | C1 | C2 | C3 | C4 | C5 | C6 | C7 | C8 |
|---|---|---|---|---|---|---|---|---|---|
| HIV | DeepRC | 0.707 | 0.798 | 0.736 | **0.834** | 0.727 | 0.666 | 0.700 | 0.711 |
|  | DeepTCR | 0.890 | 0.793 | 0.743 | 0.827 | 0.880 | 0.841 | **0.909** | 0.903 |
|  | ABMIL | **0.864** | 0.821 | 0.849 | 0.815 | 0.858 | 0.850 | 0.854 | 0.854 |
| Lupus | DeepRC | 0.478 | 0.766 | 0.832 | **0.859** | 0.433 | 0.577 | 0.514 | 0.517 |
|  | DeepTCR | 0.909 | 0.552 | 0.819 | 0.779 | 0.867 | 0.904 | **0.923** | 0.906 |
|  | ABMIL | 0.881 | 0.882 | 0.874 | 0.875 | 0.874 | 0.888 | **0.896** | 0.881 |
| COVID-19 | DeepRC | 0.659 | **0.794** | 0.751 | 0.509 | 0.701 | 0.577 | 0.532 | 0.686 |
|  | DeepTCR | 0.900 | 0.728 | 0.774 | 0.799 | 0.867 | 0.864 | **0.945** | 0.936 |
|  | ABMIL | **0.882** | 0.801 | 0.814 | 0.787 | 0.866 | 0.829 | 0.833 | 0.801 |
| Influenza | DeepRC | 0.742 | 0.714 | **0.791** | 0.664 | 0.714 | 0.655 | 0.656 | 0.684 |
|  | DeepTCR | 0.713 | 0.472 | 0.261 | 0.284 | 0.446 | 0.406 | **0.764** | 0.735 |
|  | ABMIL | 0.717 | 0.666 | 0.689 | 0.671 | 0.752 | **0.760** | 0.698 | 0.715 |
| T1D (pooled) | DeepRC | 0.869 | **0.877** | 0.875 | 0.874 | 0.861 | 0.836 | 0.581 | 0.608 |
|  | DeepTCR | 0.880 | 0.847 | 0.858 | 0.865 | 0.872 | 0.873 | **0.908** | 0.897 |
|  | ABMIL | 0.881 | 0.888 | 0.889 | 0.897 | 0.898 | 0.907 | **0.937** | 0.900 |
| TB | DeepRC | **0.676** | 0.416 | 0.644 | 0.615 | 0.597 | 0.516 | 0.456 | 0.478 |
|  | DeepTCR | 0.422 | **0.529** | 0.437 | 0.408 | 0.435 | 0.384 | 0.398 | 0.415 |
|  | ABMIL | 0.522 | 0.477 | **0.552** | 0.475 | 0.438 | 0.471 | 0.505 | 0.425 |
| RA | DeepRC | 0.562 | 0.547 | 0.605 | 0.486 | **0.614** | 0.536 | 0.512 | 0.576 |
|  | DeepTCR | 0.502 | 0.536 | 0.533 | 0.372 | 0.288 | 0.546 | **0.605** | 0.593 |
|  | ABMIL | 0.403 | 0.476 | 0.366 | 0.450 | 0.344 | **0.530** | 0.414 | 0.504 |
| CMV<sup>‡</sup> | DeepRC | 0.495 | 0.503 | 0.495 | 0.510 | **0.546** | 0.489 | 0.490 | 0.501 |
|  | DeepTCR | 0.525 | 0.517 | 0.511 | 0.513 | 0.525 | 0.521 | **0.533** | 0.531 |
|  | ABMIL | 0.526 | 0.513 | 0.519 | 0.514 | 0.521 | 0.506 | **0.530** | 0.515 |
| Rawat T1D<sup>‡</sup> | DeepRC | 0.688 | 0.620 | 0.630 | **0.709** | 0.700 | 0.707 | 0.600 | 0.643 |
|  | DeepTCR | 0.660 | 0.530 | 0.567 | 0.618 | 0.651 | 0.648 | **0.682** | 0.648 |
|  | ABMIL | 0.623 | 0.615 | 0.627 | 0.636 | 0.631 | **0.660** | 0.633 | 0.596 |

**Table B.7:** Matched-budget tuning results. For each method, the configuration selected in Stage 1 (Table B.5) is retrained with three seeds and evaluated with the same protocol as the main benchmark: pooled out-of-fold AUROC under 3-fold cross-validation, or the single held-out set for settings marked *‡* (Rawat fold 2; Emerson validation cohort). *Benchmark* is the AUROC of the default configuration reported in Table B.1; the last column is the mean *±* standard deviation across the three seeds. For the 3-fold settings, configuration selection used all three folds as validation folds (Appendix C.6), so tuned estimates may be slightly optimistic.

| Setting | Method | Selected | Benchmark | Seed 7 | Seed 21 | Seed 42 | Across seeds |
|---|---|---|---|---|---|---|---|
| HIV | DeepRC | C4 | 0.863 | 0.848 | 0.841 | 0.849 | 0.846 ± 0.004 |
|  | DeepTCR | C7 | 0.910 | 0.918 | 0.919 | 0.911 | 0.916 ± 0.004 |
|  | ABMIL | C1 | 0.865 | 0.901 | 0.914 | 0.854 | 0.890 ± 0.032 |
| Lupus | DeepRC | C4 | 0.796 | 0.859 | 0.878 | 0.864 | 0.867 ± 0.010 |
|  | DeepTCR | C7 | 0.920 | 0.941 | 0.940 | 0.933 | 0.938 ± 0.004 |
|  | ABMIL | C7 | 0.927 | 0.894 | 0.901 | 0.887 | 0.894 ± 0.007 |
| COVID-19 | DeepRC | C2 | 0.767 | 0.851 | 0.762 | 0.727 | 0.780 ± 0.064 |
|  | DeepTCR | C7 | 0.939 | 0.924 | 0.913 | 0.938 | 0.925 ± 0.012 |
|  | ABMIL | C1 | 0.809 | 0.854 | 0.889 | 0.855 | 0.866 ± 0.020 |
| Influenza | DeepRC | C3 | 0.719 | 0.762 | 0.638 | 0.783 | 0.728 ± 0.078 |
|  | DeepTCR | C7 | 0.779 | 0.829 | 0.840 | 0.819 | 0.829 ± 0.011 |
|  | ABMIL | C6 | 0.829 | 0.763 | 0.772 | 0.771 | 0.769 ± 0.005 |
| T1D (pooled) | DeepRC | C2 | 0.887 | 0.856 | 0.875 | 0.854 | 0.861 ± 0.011 |
|  | DeepTCR | C7 | 0.902 | 0.929 | 0.925 | 0.931 | 0.929 ± 0.003 |
|  | ABMIL | C7 | 0.929 | 0.936 | 0.921 | 0.925 | 0.927 ± 0.008 |
| TB | DeepRC | C1 | 0.622 | 0.600 | 0.528 | 0.550 | 0.560 ± 0.037 |
|  | DeepTCR | C2 | 0.469 | 0.454 | 0.439 | 0.456 | 0.450 ± 0.009 |
|  | ABMIL | C3 | 0.470 | 0.481 | 0.474 | 0.447 | 0.467 ± 0.018 |
| RA | DeepRC | C5 | 0.561 | 0.549 | 0.525 | 0.404 | 0.493 ± 0.078 |
|  | DeepTCR | C7 | 0.569 | 0.629 | 0.618 | 0.617 | 0.621 ± 0.007 |
|  | ABMIL | C6 | 0.528 | 0.656 | 0.633 | 0.628 | 0.639 ± 0.015 |
| CMV<sup>‡</sup> | DeepRC | C5 | 0.528 | 0.533 | 0.470 | 0.605 | 0.536 ± 0.068 |
|  | DeepTCR | C7 | 0.484 | 0.520 | 0.542 | 0.525 | 0.529 ± 0.012 |
|  | ABMIL | C7 | 0.560 | 0.564 | 0.532 | 0.546 | 0.547 ± 0.016 |
| Rawat T1D<sup>‡</sup> | DeepRC | C4 | 0.704 | 0.725 | 0.704 | 0.707 | 0.712 ± 0.011 |
|  | DeepTCR | C7 | 0.550 | 0.662 | 0.672 | 0.670 | 0.668 ± 0.006 |
|  | ABMIL | C6 | 0.679 | 0.686 | 0.715 | 0.728 | 0.710 ± 0.022 |

**Table B.8:** Pretrained sequence encoders evaluated in ABMIL. Each frozen encoder replaces the per-fold-trained CDR3 encoder; aggregation and training protocol are unchanged (Appendix C.7).

| Encoder | Checkpoint | Parameters | Embedding dim. |
|---|---|---|---|
| ESM-2 | facebook/esm2_t30_150M_UR50D | 148.4M | 640 |
| TCR-BERT | wukevin/tcr-bert | 56.8M | 768 |
| TCR-VALID | TRB 1_2_full_40 | 0.15M | 16 |

**Table B.9:**
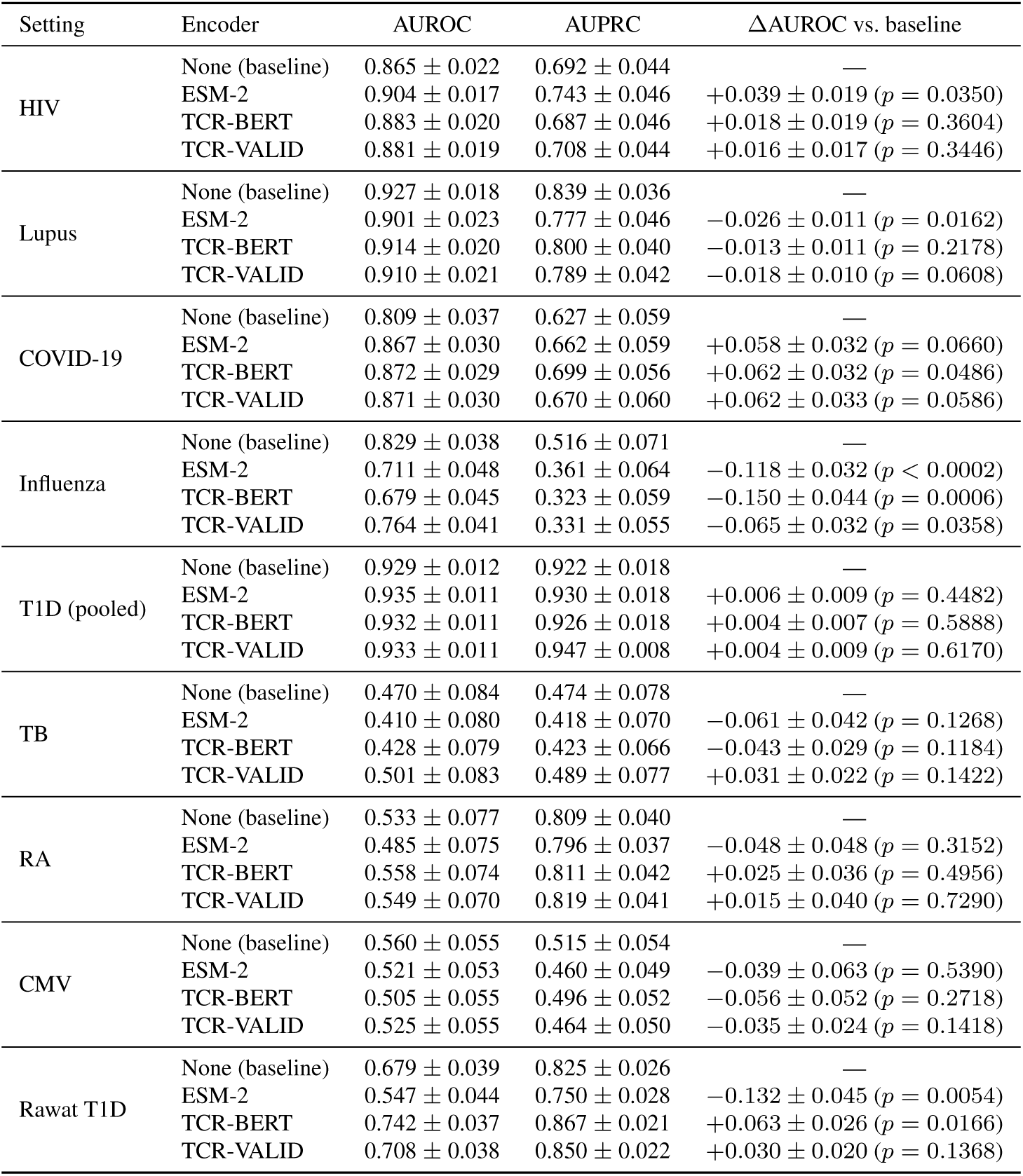
ABMIL with pretrained sequence encoders versus the baseline ABMIL with a per-fold-trained CNN encoder. AUROC and AUPRC with participant-level bootstrap standard errors; ΔAUROC is encoder minus baseline on paired resamples (*B* = 10,000), with unadjusted two-sided *p*-values. Rawat T1D and CMV use a single held-out set; all other settings use pooled 3-fold out-of-fold predictions. RA is evaluated on all 91 specimens, so its baseline differs slightly from Tables B.1 and B.2, which use the 90 specimens scored by every method.

**Table B.10:** Cross-cohort transfer of models trained on Rawat T1D, evaluated without retraining on the Mal-ID T1D cohort (92 T1D, 197 healthy/background controls) and the Mitchell T1D cohort (171 T1D, 25 healthy controls). Point estimates with participant-level bootstrap standard errors in parentheses; the best method per column is in bold. Repeated specimens are scored by their participant-mean score. Performance on the held-out Rawat fold is in Tables B.1 and B.2.

|  | Rawat → Mal-ID ($n = 289$) |  | Rawat → Mitchell ($n = 196$) |  |
|---|---|---|---|---|
| Method | AUROC | AUPRC | AUROC | AUPRC |
|---|---|---|---|---|
| Emerson et al. | **0.900** (.019) | **0.657** (.047) | 0.588 (.057) | 0.906 (.021) |
| Ostmeyer et al. | 0.404 (.035) | 0.265 (.018) | 0.389 (.059) | 0.836 (.023) |
| GIANA | 0.640 (.039) | 0.592 (.043) | 0.367 (.066) | 0.827 (.022) |
| DeepRC | 0.678 (.031) | 0.401 (.029) | 0.462 (.055) | 0.877 (.018) |
| DeepTCR | 0.499 (.034) | 0.306 (.024) | **0.645** (.055) | **0.930** (.013) |
| ABMIL | 0.438 (.032) | 0.292 (.011) | 0.482 (.059) | 0.882 (.018) |
| VJ-kmer (Reg) | 0.752 (.030) | 0.521 (.043) | 0.376 (.057) | 0.844 (.020) |
| VJ-kmer (XG) | 0.766 (.030) | 0.590 (.048) | 0.552 (.057) | 0.900 (.019) |
| Mal-ID | 0.445 (.036) | 0.283 (.020) | 0.615 (.057) | 0.921 (.014) |
| Random baseline | 0.500 | 0.318 | 0.500 | 0.872 |

**Table B.11:** Paired comparisons of each repertoire model (TCR only) against the demographics-only logistic regression on the same specimens, for the eight datasets in Figure 5c (24 tests). VJ (Reg) is the V/J-gene-usage logistic-regression sub-model of VJ-kmer (Reg). ΔAUROC is AUROC(TCR only) *−* AUROC(demographics only), with a 95% paired participant-bootstrap percentile interval (*B* = 10,000; Appendix C.5); *q*-values are Benjamini–Hochberg adjusted across the 24 tests. Positive values mean the repertoire model outperforms demographics alone. Each dataset is restricted to specimens with complete demographic covariates, so *n* can be smaller than in the main benchmark. Datasets marked *‡* are evaluated on a single held-out set (Rawat fold 2; Emerson validation cohort); all others use pooled 3-fold out-of-fold predictions.

| Dataset | Method | AUROC<br>(TCR only) | AUROC<br>(demog. only) | ΔAUROC<br>[95% CI] | *q* |
|---|---|---|---|---|---|
| HIV<br>(*n* = 253) | DeepTCR | 0.901 | 0.882 | +0.019 [−0.018, +0.058] | 0.614 |
|  | ABMIL | 0.903 | 0.882 | +0.021 [−0.017, +0.061] | 0.614 |
|  | VJ (Reg) | 0.870 | 0.882 | −0.011 [−0.062, +0.037] | 0.845 |
| Lupus<br>(*n* = 229) | DeepTCR | 0.934 | 0.809 | +0.125 [+0.069, +0.185] | 0.002 |
|  | ABMIL | 0.910 | 0.809 | +0.101 [+0.032, +0.167] | 0.018 |
|  | VJ (Reg) | 0.920 | 0.809 | +0.110 [+0.041, +0.181] | 0.006 |
| COVID-19<br>(*n* = 211) | DeepTCR | 0.896 | 0.868 | +0.028 [−0.051, +0.109] | 0.755 |
|  | ABMIL | 0.834 | 0.868 | −0.034 [−0.128, +0.062] | 0.755 |
|  | VJ (Reg) | 0.880 | 0.868 | +0.012 [−0.082, +0.106] | 0.945 |
| Influenza<br>(*n* = 231) | DeepTCR | 0.797 | 0.555 | +0.242 [+0.108, +0.373] | 0.005 |
|  | ABMIL | 0.741 | 0.555 | +0.185 [+0.056, +0.320] | 0.020 |
|  | VJ (Reg) | 0.881 | 0.555 | +0.326 [+0.209, +0.440] | 0.002 |
| TB<br>(*n* = 140) | DeepTCR | 0.423 | 0.464 | −0.041 [−0.163, +0.087] | 0.755 |
|  | ABMIL | 0.470 | 0.464 | +0.006 [−0.196, +0.210] | 0.973 |
|  | VJ (Reg) | 0.472 | 0.464 | +0.008 [−0.185, +0.204] | 0.973 |
| RA<br>(*n* = 91) | DeepTCR | 0.453 | 0.528 | −0.075 [−0.325, +0.171] | 0.777 |
|  | ABMIL | 0.660 | 0.528 | +0.132 [−0.049, +0.308] | 0.367 |
|  | VJ (Reg) | 0.532 | 0.528 | +0.004 [−0.199, +0.209] | 0.973 |
| CMV<sup>‡</sup><br>(*n* = 120) | DeepTCR | 0.501 | 0.560 | −0.059 [−0.195, +0.083] | 0.755 |
|  | ABMIL | 0.526 | 0.560 | −0.034 [−0.183, +0.111] | 0.845 |
|  | VJ (Reg) | 0.584 | 0.560 | +0.024 [−0.112, +0.160] | 0.872 |
| Rawat T1D<sup>‡</sup><br>(*n* = 204) | DeepTCR | 0.613 | 0.819 | −0.206 [−0.294, −0.120] | 0.002 |
|  | ABMIL | 0.679 | 0.819 | −0.140 [−0.223, −0.060] | 0.002 |
|  | VJ (Reg) | 0.720 | 0.819 | −0.099 [−0.182, −0.020] | 0.031 |

**Table B.12:** Participant-level demographics-only baselines for a subset of cohorts. Logistic regression on the listed covariates alone, evaluated on participants with complete covariates; AUROC and AUPRC with participant-level bootstrap standard errors. Rawat T1D and CMV use the held-out evaluation set (Rawat fold 2; Emerson validation cohort); RA and TB use pooled 3-fold out-of-fold predictions at the participant level. Ancestry is not recorded for Rawat, RA, or TB. Because RA and TB include multiple specimens per participant, these participant-level baselines differ from the specimen-level demographics-only values in Table B.11, which are fit and scored per specimen to match the repertoire models. Rawat values differ from Table B.11 because all three Rawat models here are trained and evaluated only on participants with a recorded HLA-risk category (194 of 204 in the held-out set). HLA-risk category is the HLA-DR genotype group recorded in the Rawat metadata, defined by the T1D risk haplotypes DR3 and DR4 (DR3/3, DR3/4, DR4/4, DR3/X, DR4/X, or DRX/X, where X is neither DR3 nor DR4), and is one-hot encoded.

| Cohort | Features | *n* (cases/controls) | AUROC | AUPRC |
|---|---|---|---|---|
| TB | Random baseline | 67 (27/40) | 0.500 | 0.403 |
|  | Age |  | 0.505 ± 0.072 | 0.420 ± 0.057 |
|  | Age + sex |  | 0.497 ± 0.072 | 0.411 ± 0.057 |
|  | Age + sex + recruitment cohort |  | 0.539 ± 0.074 | 0.481 ± 0.071 |
| RA | Random baseline | 84 (64/20) | 0.500 | 0.762 |
|  | Age |  | 0.406 ± 0.063 | 0.748 ± 0.040 |
|  | Age + sex |  | 0.419 ± 0.064 | 0.753 ± 0.040 |
| CMV | Random baseline | 120 (51/69) | 0.500 | 0.425 |
|  | Age |  | 0.522 ± 0.053 | 0.443 ± 0.043 |
|  | Age + sex |  | 0.521 ± 0.054 | 0.444 ± 0.044 |
|  | Age + sex + race/ethnicity |  | 0.560 ± 0.055 | 0.560 ± 0.053 |
| Rawat T1D | Random baseline | 194 (135/59) | 0.500 | 0.696 |
|  | Age |  | 0.824 ± 0.031 | 0.905 ± 0.027 |
|  | Age + sex |  | 0.824 ± 0.031 | 0.905 ± 0.027 |
|  | Age + sex + HLA-risk category |  | 0.866 ± 0.029 | 0.921 ± 0.026 |

**Table B.13:** AUROC (**Top**) / AUPRC (**Bottom**) across *k*-mer featurizations of the VJ-kmer logistic-regression model (VJ-kmer (Reg)) on five disease classification tasks. We sweep *k*-mer length (*k ∈ {*3, 4, 5*}*) and gapping (ungapped vs. gapped+ungapped combined) for two model variants: the full *Ensemble* that combines *k*-mer and V/J-gene features, and a *k-mer-only* ablation. The *V/J only* baseline uses V/J-gene features alone and is therefore independent of the *k*-mer sweep. We also report the per-disease range Δ = max *−* min across the configurations. T1D refers to the Mal-ID T1D cohort alone (96 T1D versus 197 healthy controls). **Take-away:** on every disease the ensemble varies less with the *k*-mer featurization than the *k*-mer-only model (Δ AUROC 0.010–0.061 versus 0.023–0.401), most markedly on Influenza, where the *k*-mer-only model spans 0.457–0.859 and the ensemble 0.857–0.917, indicating that V/J features absorb most of the variation introduced by the *k*-mer choice.

| Model | Featurization | HIV | Lupus | COVID-19 | Influenza | T1D |
|---|---|---|---|---|---|---|
| Ensemble<br>(*k*-mer + V/J) | *k* = 3, ungapped | 0.9149 | 0.9261 | 0.9064 | 0.8569 | 0.9599 |
|  | *k* = 3, gapped+ungapped | 0.9026 | 0.9265 | 0.9094 | 0.8660 | 0.9638 |
|  | *k* = 4, ungapped | 0.8730 | 0.9321 | 0.9082 | 0.9085 | 0.9788 |
|  | *k* = 4, gapped+ungapped | 0.8700 | 0.9309 | 0.9220 | 0.8730 | 0.9808 |
|  | *k* = 5, ungapped | 0.8733 | 0.9219 | 0.8750 | 0.9174 | 0.9798 |
|  | *k* = 5, gapped+ungapped | 0.8969 | 0.9217 | 0.9064 | 0.8746 | 0.9765 |
|  | Range (Δ) | **0.0449** | **0.0104** | **0.0470** | **0.0605** | **0.0209** |
| *k*-mer only | *k* = 3, ungapped | 0.8926 | 0.9104 | 0.8130 | 0.6126 | 0.9423 |
|  | *k* = 3, gapped+ungapped | 0.8844 | 0.9122 | 0.8103 | 0.4571 | 0.9527 |
|  | *k* = 4, ungapped | 0.7694 | 0.9026 | 0.8270 | 0.7484 | 0.9857 |
|  | *k* = 4, gapped+ungapped | 0.7931 | 0.9135 | 0.8148 | 0.7458 | 0.9824 |
|  | *k* = 5, ungapped | 0.8282 | 0.9256 | 0.8778 | 0.8426 | 0.9741 |
|  | *k* = 5, gapped+ungapped | 0.8200 | 0.9231 | 0.8869 | 0.8586 | 0.9798 |
|  | Range (Δ) | 0.1231 | 0.0230 | 0.0767 | 0.4014 | 0.0435 |
| V/J only | — | 0.8601 | 0.9282 | 0.8575 | 0.8763 | 0.9249 |
| Model | Featurization | HIV | Lupus | COVID-19 | Influenza | T1D |
| Ensemble<br>(*k*-mer + V/J) | *k* = 3, ungapped | 0.8101 | 0.7656 | 0.7625 | 0.5981 | 0.8928 |
|  | *k* = 3, gapped+ungapped | 0.8318 | 0.7653 | 0.7645 | 0.6427 | 0.8891 |
|  | *k* = 4, ungapped | 0.7508 | 0.7807 | 0.7594 | 0.6727 | 0.9245 |
|  | *k* = 4, gapped+ungapped | 0.7552 | 0.7788 | 0.7728 | 0.6354 | 0.9318 |
|  | *k* = 5, ungapped | 0.7323 | 0.7850 | 0.7459 | 0.7403 | 0.9456 |
|  | *k* = 5, gapped+ungapped | 0.8020 | 0.7868 | 0.7537 | 0.6596 | 0.9352 |
|  | Range (Δ) | **0.0995** | **0.0216** | **0.0269** | **0.1422** | **0.0565** |
| *k*-mer only | *k* = 3, ungapped | 0.8398 | 0.7113 | 0.5685 | 0.2176 | 0.8728 |
|  | *k* = 3, gapped+ungapped | 0.8306 | 0.7124 | 0.5914 | 0.1505 | 0.8615 |
|  | *k* = 4, ungapped | 0.6843 | 0.7292 | 0.7038 | 0.4043 | 0.9548 |
|  | *k* = 4, gapped+ungapped | 0.7200 | 0.7594 | 0.6543 | 0.3834 | 0.9398 |
|  | *k* = 5, ungapped | 0.7375 | 0.8531 | 0.7547 | 0.5528 | 0.9119 |
|  | *k* = 5, gapped+ungapped | 0.7758 | 0.8329 | 0.7665 | 0.6120 | 0.9485 |
|  | Range (Δ) | 0.1555 | 0.1418 | 0.1980 | 0.4616 | 0.0933 |
| V/J only | — | 0.7130 | 0.7732 | 0.7500 | 0.6355 | 0.7727 |

**Table B.14:** CLI arguments used for the Mal-ID runs.

| Component | CLI argument | Value | Notes |
|---|---|---|---|
| Model 1 | -model1-n-pcs | 15 |  |
| Model 1 | -model1-l1-ratio | per-locus | automatically sets 1.0 (lasso) for TCR |
| Model 1 | -model1-model-name | lasso_cv |  |
| Model 2 | -model2-sequence-identity-threshold | per-locus | automatically sets 0.90 for TCR |
| Model 2 | -model2-p-values | [0.0005, 0.001, 0.005, 0.01, 0.05] |  |
| Model 2 | -model2-retrain-on-full-train | False |  |
| Model 2 | -model2-abstention-strategy | fill_models13_mean |  |
| Model 3 | -model3-aggregation-strategy | entropy_percentile_cutoff |  |
| Model 3 | -model3-entropy-bottom-percentile | 0.01 |  |
| Model 3 | -model3-tuning-cv-splits | 3 |  |
| Model 3 | -model3-n-estimators-stage2 | 100 |  |
| Ensemble | -model2-abstention-strategy | fill_models13_mean |  |
| Caching | -clone-id-use-aa |  | immunoSEQ datasets only |
| Embeddings | -batch-size | 15,000 |  |

**Table B.15:** Handling model-2 abstentions in the Mal-ID ensemble. We compare the original abstention-based ensemble to two strategies for imputing missing model-2 outputs on abstained specimens: replacing the missing value with 0.5, or replacing it with the per-specimen mean of that specimen’s model-1 and model-3 predictions. We report accuracy, AUROC, and AUPRC across diseases, together with abstention rates and the numbers of abstained train and test specimens. For the abstention-based ensemble, abstained specimens are counted as erroneous predictions for accuracy and excluded from AUROC and AUPRC computation.

|  | Accuracy |  |  | AUROC |  |  | AUPRC |  |  | Abstention |  |  |
|---|---|---|---|---|---|---|---|---|---|---|---|---|
| Disease | M2-abstained | M2-fill-0.5 | M2-fill-mean-M1-M3 | M2-abstained | M2-fill-0.5 | M2-fill-mean-M1-M3 | M2-abstained | M2-fill-0.5 | M2-fill-mean-M1-M3 | Rate | Test (#) | Train (#) |
| HIV | 0.8475 | 0.8678 | 0.8678 | 0.9131 | 0.9155 | 0.9160 | 0.7308 | 0.7442 | 0.7470 | 2.0% | 6 | 0 |
| Lupus | 0.8276 | 0.8851 | 0.8851 | 0.9159 | 0.9446 | 0.9450 | 0.7910 | 0.8630 | 0.8626 | 5.0% | 13 | 5 |
| COVID-19 | 0.9412 | 0.9451 | 0.9451 | 0.9859 | 0.9857 | 0.9858 | 0.9386 | 0.9384 | 0.9387 | 0.4% | 1 | 0 |
| Influenza | 0.8761 | 0.9444 | 0.9444 | 0.9751 | 0.9691 | 0.9745 | 0.9535 | 0.9190 | 0.9427 | 7.3% | 17 | 7 |
| T1D | 0.9215 | 0.9386 | 0.9386 | 0.9819 | 0.9660 | 0.9633 | 0.9551 | 0.9403 | 0.9383 | 2.7% | 8 | 1 |

## C Technical Appendices

### C.1 Dataset Preprocessing

All repertoire files consumed by the benchmarked methods are harmonized to a single AIRR-compliant schema with cdr3_aa, v_call, and j_call as the sequence-level columns, regardless of which source they originated from. The Mal-ID and immunoSEQ data enter the pipeline in different raw formats and therefore follow different but conceptually parallel cleanup paths, described below.

#### Mal-ID

The Mal-ID release is distributed in an internal, per-participant tabular format produced by an IgBLAST-based annotation pipeline. For each participant file we (i) drop non-productive rearrangements and low-confidence V assignments, (ii) strip the whitespace inserted by IgBLAST between residues in each amino-acid field and uppercase the result, (iii) drop any row with a missing CDR3, V, or J call or with a non-standard amino-acid character in its CDR3 (e.g., * for a stop codon, X for an ambiguous residue), and (iv) collapse a small number of V-gene alleles that are known to be indistinguishable under the FR3 primers used in the Mal-ID assay to a canonical allele, following Omer et al. [5]. The cleaned per-participant tables are renamed into AIRR column conventions and split by repertoire_id into one file per specimen, yielding the 550 specimen-level files.

#### immunoSEQ cohorts

The five immunoSEQ cohorts (Mitchell T1D, TB, RA, Rawat T1D, and Emerson CMV) are distributed in Adaptive immunoSEQ tabular formats, which differ from AIRR in column names and gene nomenclature. We map the CDR3 amino-acid, V-gene, and J-gene columns (e.g., aminoAcid, vGeneName, jGeneName) to cdr3_aa, v_call, and j_call. Gene names are converted from Adaptive to IMGT nomenclature (e.g., TCRBV07-02 to TRBV7-2); for ambiguous calls listing several genes, we keep the first, and we strip allele annotations to match the gene-level resolution of Mal-ID. We also remove the first and last CDR3 residue (the conserved flanking cysteine and phenylalanine), so that the CDR3 definition matches the Mal-ID files. We retain only productive, in-frame rearrangements, using each source’s productivity annotation (sequenceStatus or frame_type equal to In), and remove rows with a missing or unresolved CDR3, V, or J call, a non-standard amino acid in the CDR3, or a V/J assignment without gene-level resolution. No V-assignment score threshold is applied. Finally, we exclude any specimen with fewer than 1,000 productive rearrangements, since such repertoires carry too little signal to be informative and are dominated by sampling noise. This threshold is applied before the gene-resolution and CDR3 filters above, so a retained specimen can fall slightly below 1,000 sequences after preprocessing (e.g., the RA synovial-fluid sample RA6-SF, with 904); such specimens are excluded by Mal-ID’s own minimum-size filter, which is why Mal-ID scores 90 of the 91 RA specimens (Table B.1). From Rawat cohort 1 (1,393 repertoires), we retain 426 participants with T1D and 188 unrelated, islet-autoantibody-negative controls, excluding 625 first-degree relatives, 59 second-degree relatives, and 95 autoantibody-positive participants without diabetes. For Emerson CMV, the 761 specimens with known serostatus (641 discovery, 120 validation) are eligible for classification; 25 discovery-cohort specimens with unknown status are kept in the released metadata but excluded from classification, and one discovery-cohort specimen falls below the 1,000-sequence threshold (Table B.3).

#### Cross-source gene-label reconciliation

The Mal-ID and immunoSEQ pipelines independently produce AIRR-style V-gene labels, but the two conventions disagree on single-member TRBV families: Mal-ID (IMGT) writes these without a trailing -1, while the Adaptive-derived files retain it (e.g., TRBV13 vs. TRBV13-1). During preprocessing of every immunoSEQ cohort, we collapse the Adaptive-style -1 suffix on the ten singleton families (TRBV2, TRBV9, TRBV13–TRBV15, TRBV18, TRBV19, TRBV27, TRBV28, TRBV30), so that the same gene receives the same label in all sources. All released repertoires therefore share a single labeling convention, and within-cohort, pooled, and zero-shot cross-cohort evaluations (e.g., Rawat-trained models applied to Mal-ID) see identical V-gene labels.

#### Conversion to immuneML-compatible AIRR format

We additionally export the cleaned repertoires into a common AIRR Rearrangement schema for immuneML [30], preserving every row and its original cdr3_aa and V/J calls. We standardize locus and Boolean fields, supply sequence and repertoire identifiers, and map available counts to duplicate_count, leaving unavailable values empty. Junction sequences are reconstructed from source annotations or, for Rawat and Emerson, by locating cdr3_aa in three-frame translations of the nucleotide sequence and extracting one flanking codon on each side. Reconstructed junctions must have valid nucleotide content and length and translate consistently with the recorded CDR3; otherwise, junction fields remain empty and the row is retained. Comma-separated metadata files preserve specimen and participant identifiers, disease labels, available demographics, and predefined folds. We also provide setting-specific training and test metadata files and example immuneML configurations that use these splits. The configurations specify region_type: IMGT_CDR3, so immuneML analyzes the same cdr3_aa amino-acid sequences used in BenchRep-T.

### C.2 Methods Details

#### C.2.1 Statistical Disease-Signature Methods

##### Emerson et al. (2017)

The immunosequencing model of Emerson et al. [7] identifies individual TCR*β* clones that are differentially present in cases versus controls and uses their occurrence counts as a generative signal. For each unique (V gene, J gene, CDR3 amino-acid sequence) triplet observed in at least two training subjects, we perform a one-sided Fisher’s exact test on the 2*×*2 contingency table of presence/absence against disease label. Triplets with *p*-value below a threshold form the diagnostic set *T* . For each subject *i* we compute the repertoire size *n_i_* (number of unique triplets in the subject) and the diagnostic burden *k_i_* (number of those triplets that fall in *T*), and fit class-conditional Beta-Binomial distributions BB(*k_i_ | n_i_, α_c_, β_c_*) for *c ∈ {*disease, healthy*}* by minimizing the negative log-likelihood. At test time we compute log-posterior odds with a Laplace-smoothed class prior (*N_c_* + 1)*/*(*N* + 2). The model exposes a single core hyperparameter, the Fisher’s exact test *p*-value threshold used to select disease-associated diagnostic TCRs. It is selected on an internal 90/10 train/validation split by validation AUROC (Appendix C.4), with 10*^−^*^4^, the cross-validation-optimized value of the original paper, as the default.

##### Ostmeyer et al. (2019)

Ostmeyer et al. [24] models a repertoire as a bag of 4-mer motifs. After stripping the first three and last three residues of each CDR3, every overlapping 4-mer is encoded as a 20-dimensional vector by concatenating the five Atchley factors [40] of its residues, augmented with the log of its clone-size-weighted relative abundance to give a 21-dimensional feature. A logistic regression is trained on these bags under a max-aggregation MIL objective, so each bag’s gradient flows only through its highest-scoring “witness” 4-mer. We follow the authors’ prescription of repeated solver restarts with Gaussian weight initialization and no L1/L2 regularization, keeping the restart with the lowest training loss. Optimization is controlled by two parameters: the number of Gaussian restarts, i.e., the number of random L-BFGS-B initializations from which the lowest-training-loss solution is retained (we use 200 as a computational approximation to the original 100,000–375,000; Appendix C.4), and the maximum number of L-BFGS-B iterations (default 1000), a per-restart cap that serves as a safety bound since runs typically converge within 50–200 gradient evaluations.

### C.2.2 Ensemble of Engineered Repertoire-level Representations

#### V/J gene + Gapped *k*-mer (VJ-kmer) ensemble

We represent each repertoire using two simple feature dictionaries of relative frequencies: (i) V-gene and J-gene usage, each normalized within its own gene type, and (ii) all CDR3 4-mers plus their four single-position gapped variants (“X_YZ”, etc.), normalized by the total *k*-mer count. Each feature set is passed to an *L*_1_-penalized logistic regression; the inverse-regularization strength *C* is selected independently for each sub-model by 5-fold stratified CV using AUROC, from [0.001, 0.003, 0.01, 0.03, 0.1] for the *k*-mer sub-model and [1.0, 0.2, 0.1, 0.05, 0.03] for the V/J sub-model. The two sub-model probabilities are combined as *p* = *α · p*_kmer_ + (1 *− α*) *· p*_VJ_, with *α ∈ {*0.0, 0.1*, . . . ,* 1.0*}* tuned on an internal 80*/*20 validation split. We report the performance of two sub-models (with *k*-mer features only and V/J gene features only) in isolation as ablations (Table B.13).

We additionally evaluate an XGBoost [36] variant that swaps each *L*_1_-penalized logistic regression base learner for a gradient-boosted tree classifier, keeping the same feature dictionaries and the same *α* sweep. Here we use binary:logistic objective with up to 1,000 boosting rounds and early stopping after 20 rounds without improvement on a validation set. Hyperparameter tuning is conducted through a two-stage deterministic grid search:

1. Stage 1 sweeps max_depth *∈ {*3, 4, 5, 6*} ×* learning_rate *∈ {*0.01, 0.03, 0.05, 0.1*}* (16 combinations).
2. Stage 2 fixes the best Stage 1 values, then sweeps subsample *∈ {*0.7, 0.8, 1.0*} ×* colsample_bytree *∈ {*0.7, 0.8, 1.0*} ×* min_child_weight *∈ {*1, 3, 5*}* (27 combinations).

#### Mal-ID, Zaslavsky et al. (2025)

Mal-ID (MAchine Learning for Immunological Diagnosis) [12] is an ensemble framework that combines three complementary repertoire representations for the BCR and TCR repertoires separately. These include (i) a repertoire-composition model based on normalized V/J gene usage and, for BCR, somatic hypermutation features; (ii) a convergent-clustering model that groups sequences by V gene, J gene, and CDR3 length, clusters sequences within each group by CDR3 amino acid similarity, identifies disease-enriched clusters by statistical testing, and trains a classifier on cluster-count features; and (iii) a protein language model pipeline that embeds individual CDR3 sequences with ESM-2 [37], scores them with sequence-level classifiers specialized by V gene, and aggregates these predictions to the specimen level with a second-stage classifier. Further details regarding adaptation applied in this study are in Appendix C.3. The Mal-ID re-implementation is available at https://github.com/liel-cohen/Mal-ID-Lite.

Mal-ID-Lite arguments used for the study runs are listed in Table B.14.

### C.2.3 Clustering-based Methods

#### GIANA, Zhang et al. (2021)

GIANA [28] uses a precomputed substitution-matrix embedding, derived from BLOSUM-based distances and multi-dimensional scaling, to encode each CDR3 into a 96-dimensional isometric vector. Training sequences are clustered with a length-adjusted isometric-distance threshold, followed by V-gene and Smith-Waterman refinement. Reference clusters are built from training sequences only and held fixed while test sequences are assigned to them in query mode, so test repertoires never influence cluster membership or disease scores. For each reference cluster, we compute the fraction of training sequences from the target disease, excluding clusters with more than 100 reference TCRs. A test specimen’s score is the mean disease fraction across its TCRs that are assigned to reference clusters. TCRs without a reference cluster assignment are omitted from the mean, and specimens with no assigned TCRs receive score zero. Up to 10,000 sequences per specimen are kept. We use method defaults for parameters, including isometric distance threshold 7 in exact mode, Smith-Waterman score threshold 3.3, and V-gene similarity threshold 3.7.

### C.2.4 Deep Learning-based Methods

#### Attention-based Multiple Instance Learning (ABMIL)

We implement an attention-based MIL model [27] that replaces hand-designed features with end-to-end learned embeddings. A learned amino-acid embedding (hidden dim. 64) is passed through a three-layer 1-D CNN (kernel size 5, channel dimensions 32 *→* 64 *→* 128, LeakyReLU activations, dropout 0.25) with global max-pooling to produce a per-sequence vector; learned V- and J-gene embeddings (hidden dim. 48 each) are concatenated to this vector. A gated-attention aggregator pools per-sequence features into a repertoire-level representation via weighted summation, followed by a single linear layer and sigmoid classifier. The model is trained with Adam (learning rate 5*×*10*^−^*^4^, weight decay 10*^−^*^4^) to minimize binary cross-entropy with positive-class weighting (*n*_neg_*/n*_pos_), using early stopping (patience 10 epochs) on an internal 20% validation split of bags. Training runs for up to 100 epochs; each epoch subsamples at most 10,000 sequences per bag.

#### DeepRC, Widrich et al. (2020)

DeepRC [19] is a multiple instance repertoire classifier based on modern Hopfield networks. Each CDR3 is represented by one-hot amino-acid features (20 channels) augmented with three positional features. A single-layer 1-D convolutional sequence encoder (Conv1d with 23 input channels, 32 kernels of size 9, SELU activation, and global max-pooling) maps each sequence to a fixed-dimensional embedding. We use the authors’ PyTorch implementation with parameter defaults retained from their CNN sequence encoder and attention architecture. Training uses Adam (learning rate 10*^−^*^4^, no *L*_2_ weight decay) to minimize binary cross-entropy over a budget of 10,000 gradient updates with a batch size of 32 repertoires. Up to 10,000 sequences are subsampled per repertoire during training.

#### DeepTCR, Sidhom et al. (2021)

DeepTCR [26] is a convolutional multi-instance learning architecture for repertoire-level prediction. Each CDR3 is represented with a trainable amino-acid embedding (hidden dim. 64) passed through three 1-D convolutional layers (widths 12, 32, and 64; kernel size 5; LeakyReLU activation; dropout 0 by default), while V- and J-gene identities are captured by learned gene embeddings (hidden dim. 48 each). Sequences sharing identical CDR3s are first aggregated by summing their clone counts; the model then maps each unique sequence to a learned space of 64 concepts and aggregates across the repertoire via a frequency-weighted average. The model is implemented in TensorFlow and trained with Adam (lr=10*^−^*^3^) to minimize softmax cross-entropy augmented with a per-sample hinge regularization term (max(0*, ℓ −* 0.1)) using a batch size of 25 repertoires. Early stopping monitors validation loss over a rolling window of 10 epochs and halts training when relative improvement falls below a threshold of 0.25, subject to a minimum of 25 epochs.

#### Embedding provenance

ABMIL and DeepTCR use randomly initialized amino-acid (64-dimensional) and V/J-gene (48-dimensional each) embeddings trained per fold. DeepRC uses no lookup embeddings: 20 one-hot amino-acid channels plus 3 fixed positional features. Ostmeyer et al. (Atchley factors) and GIANA (BLOSUM-derived) use fixed, non-learned representations. ESM-2 in Mal-ID model 3 is the only pretrained representation among the nine benchmarked methods.

### C.3 Mal-ID Adaptations

In the original formulation, the three BCR and three TCR base models are combined by a logistic-regression metamodel to predict immune status. Mal-ID was introduced as a multiclass framework operating jointly on BCR and TCR repertoires, with the strongest performance obtained by combining both cell types and all three model components. To integrate Mal-ID into BenchRep-T, we implemented a streamlined reimplementation of the published pipeline, which preserves the model of the original method while making it easier to run consistently across benchmark tasks, folds, and future reuse settings. Two adjustments were introduced for the BenchRep-T benchmark setting.

#### Adaptive entropy threshold in model 3

In the published Mal-ID configuration, model 3 aggregates sequence-level class probabilities after filtering out high-entropy sequences, so that specimen-level features are computed from lower-entropy, more class-informative predictions. In our benchmark setting, we found that using a fixed entropy cutoff could, in some folds, filter all sequences from a specimen or even nearly all sequences in the training data for a given model configuration, leading to degenerate post-aggregation features. To avoid this behavior while preserving the intended preference for lower-entropy sequences, we replaced the fixed cutoff with a training-set-based percentile threshold (0.01%). For each training fold, we estimate the entropy distribution of sequence-level predictions on the training data and store the corresponding 0.01th-percentile value as part of the fitted model. This learned cutoff is then applied unchanged at inference time to any validation or test repertoire scored by that model. The result is a filter that adapts to the training-time score distribution while avoiding collapse of the post-filtering sequence set.

#### Replacing model-2 abstentions in the ensemble

In the original Mal-ID pipeline, model 2 abstains when no sequence in a specimen matches any disease-enriched cluster, and these abstentions propagate to the ensemble model, which abstains from them as well. In BenchRep-T, however, such abstentions would make AUROC and AUPRC comparisons across methods difficult, because different methods would effectively be evaluated on different subsets of specimens. We therefore replaced missing model-2 outputs with filled values in both the ensemble training and test matrices.

We compared three strategies: retaining abstentions, filling model 2 abstained sample predictions with a constant value of 0.5, and filling with the mean of the corresponding model-1 and model-3 predictions. As shown in Table B.15, the two filling strategies produced very similar results overall, with identical accuracy, and the model-1/model-3 mean fill was slightly better on average (mean AUROC 0.957 vs. 0.956; mean AUPRC 0.886 vs. 0.881). Relative to retaining abstentions, filling increased accuracy on every disease, since abstained specimens otherwise count as errors, and increased AUROC and AUPRC on HIV and Lupus, left them nearly unchanged on COVID-19, and decreased them on Influenza and T1D (e.g., T1D AUROC 0.982 to 0.963). The abstention-based AUROC and AUPRC are computed only on non-abstained specimens and are therefore not directly comparable to the filled metrics. Filling enabled every specimen to receive an ensemble prediction, so we use the model-1/model-3 mean fill strategy throughout BenchRep-T.

### C.4 Reproduction fidelity

This section covers the methods that we reimplemented or adapted from prior work, and checks each against its published description or results. ABMIL and both VJ-kmer variants are new to this work and have no published reference.

#### C.4.1 Emerson et al

We reimplemented the published method [7]: (V gene, J gene, CDR3) clonotypes enriched in cases by a one-sided Fisher’s exact test form the diagnostic set, class-conditional Beta-Binomial distributions are fit to each subject’s diagnostic burden given repertoire size, and test subjects are scored by log-posterior odds. The BenchRep-T CMV setting follows the original study design, training on the discovery cohort and evaluating on the independent 120-subject validation cohort, so it doubles as a reproduction check. Our reimplementation reaches AUROC 0.893 (95% participant-bootstrap CI 0.826–0.950) on this cohort; the 0.94 reported by Emerson et al. lies within this interval, and the reimplementation remains the strongest method on this setting by a wide margin (Table B.1). Two differences from the original protocol may contribute to the remaining gap: the *p*-value threshold is selected on an internal 90/10 train/validation split rather than by leave-one-out cross-validation, and repertoires pass through the harmonized BenchRep-T preprocessing (Appendix C.1), including gene-level V/J calls.

#### C.4.2 Ostmeyer et al

We reproduced the published representation (4-mers encoded by Atchley factors, with a relative-abundance feature), the max-aggregation multiple instance scoring, and the random-restart L-BFGS-B optimization [24]. The restart count is a computational approximation rather than an exact reproduction. The original did not use 250,000 restarts uniformly: its Table 3 reports 100,000–375,000 across model variants. At our measured cost of 13.2 CPU-hours for one 200-restart fold, 250,000 restarts would require about 16,400 CPU-hours (*∼*685 days) per fold. We observe no considerable performance differences across restart counts, and we use 200 restarts throughout.

#### C.4.3 GIANA

GIANA uses the author-provided GIANA 4.1 encoding and clustering routines within its published reference/query framework [28]. Our only change is to the query orchestration for cross-validation: reference clusters are built from training data only and held fixed during test assignment, so test data cannot alter the training-derived clusters (Appendix C.2). The same pipeline performs well on Influenza, Lupus, and T1D, so its below-chance AUROC on RA and TB reflects a mismatch between its inductive bias and these phenotypes rather than an implementation fault. GIANA assumes that cases share convergent, disease-associated TCR clusters, and scores a repertoire in a fixed direction by the fraction of neighboring reference TCRs contributed by cases. Methods that learn flexible discriminative features collapse toward chance (AUROC *≈*0.5) when a cohort carries little stable signal. GIANA instead assigns controls higher scores whenever control repertoires are more public and convergent than those of RA patients or TB progressors, which yields AUROCs well below 0.5.

### C.4.4 DeepRC

We ran our adaptation of the authors’ code on the 5-fold splits of the Emerson CMV discovery cohort released by Widrich et al. [19], under the method’s own nested cross-validation and preprocessing. It reproduces the published result: AUROC 0.8271 *±* 0.0353 over five folds, versus the reported 0.831 *±* 0.002. This setting differs from the BenchRep-T CMV task, in which models are trained on the discovery cohort and evaluated on the independent 120-specimen validation cohort (AUROC 0.528 for DeepRC), which characterizes cross-cohort transfer.

#### C.4.5 DeepTCR

DeepTCR is run with the authors’ released implementation [26], unmodified apart from loading the benchmark inputs, so the model, training procedure, and inference are those of the original. We therefore did not perform a separate reproduction.

#### C.4.6 Mal-ID

We evaluated Mal-ID-Lite on the original Mal-ID data in the original setting (multiclass, TCR only), following the source protocol of pooled accuracy and per-fold probability-based metrics. The full TCR metamodel closely reproduced every TCR quantity reported in Table S2 of Zaslavsky et al. [12]: accuracy 75.5% (versus 75.1%), one-versus-one AUROC 0.955 *±* 0.005 (versus 0.952 *±* 0.003), AUPRC 0.947*±*0.001 (versus 0.943*±*0.008), and one abstaining specimen out of 550 (0.18%, versus 0.4%). Model 1 also matched closely, whereas models 2 and 3 deviated in opposite directions by roughly twice the fold-level standard deviation reported in the original. Two factors explain this. First, the per-component rows of Table S2 are ensemble metamodels refit on feature subsets rather than standalone base-model evaluations, so they are not directly comparable to our base models. Second, the original assignments of samples to the inner train/validation partitions and to the base-model cross-validation splits were not released, and base-model selection and training are sensitive to these assignments at the available class sizes. The metamodel, which operates on a low-dimensional feature matrix, is correspondingly more stable, consistent with what we observe.

### C.5 Uncertainty estimation and significance testing

All uncertainty estimates are computed from fixed out-of-fold predictions, with no retraining. For each cohort, we draw *B* = 10,000 bootstrap resamples of participants with replacement within each class stratum, so that every resample keeps the original numbers of cases and controls; repeated specimens from the same participant are resampled together. The index sets are drawn once per cohort and reused for every method, so estimates are paired across methods. We report the standard deviation of the replicate metrics as the bootstrap standard error and the 2.5th and 97.5th percentiles as the 95% confidence interval. A result is above chance when the lower bound of its 95% interval exceeds the random baseline (0.5 for AUROC; the positive-class rate for AUPRC). Pairwise method comparisons use both the paired bootstrap and DeLong’s test, with Benjamini–Hochberg (BH) correction within each family of comparisons.

#### Paired comparisons

For two methods *M*_1_ and *M*_2_ scored on the same specimens, the paired bootstrap evaluates both methods on each of the *B* shared participant resamples and forms the replicate differences Δ*_b_* = AUROC*_b_*(*M*_1_)*−* AUROC*_b_*(*M*_2_). We report the full-sample difference ΔAUROC, the standard deviation of Δ*_b_* as its standard error, and the 2.5th and 97.5th percentiles of Δ*_b_* as its 95% interval. The two-sided *p*-value inverts the percentile interval, *p* = min*{*1, 2 min(*p_−_, p*_+_)*}* with *p_−_* = (#*{*Δ*_b_ ≤* 0*}* + 1)*/*(*B* + 1) and *p*_+_ = (#*{*Δ*_b_ ≥* 0*}* + 1)*/*(*B* + 1), so *p <* 0.05 exactly when the 95% interval excludes zero; its smallest attainable value is 2*/*(*B* + 1) *≈* 2 *×* 10*^−^*^4^. Because resampling is by participant, the test respects repeated specimens from the same donor. As a cross-check we also apply DeLong’s test for two correlated AUROCs, which has a closed form but treats specimens as independent and is therefore anti-conservative when a participant contributes several specimens (as in TB, with 140 specimens from 67 participants).

We test two families. *Ensemble versus deep learning* (Table B.4) compares each feature-ensemble method (VJ-kmer (XG), Mal-ID) with each deep-learning method (DeepRC, DeepTCR) on all nine settings (36 tests), with BH correction applied across the 36 tests separately for the bootstrap and DeLong *p*-values. *Repertoire versus demographics* (Table B.11) compares each repertoire-only model (DeepTCR; ABMIL; VJ (Reg), the V/J sub-model of VJ-kmer (Reg)) with the demographics-only logistic regression on the eight datasets of Figure 5c (24 tests, BH-corrected across the 24). For this family both models are scored on the same complete-covariate specimen set of each dataset (Appendix C.13), and only the paired bootstrap is reported.

### C.6 Matched-budget hyperparameter tuning

To test whether the deep learning methods were disadvantaged by their default hyperparameters, we tuned ABMIL, DeepTCR, and DeepRC under a matched budget: each method receives eight configurations to choose from (Table B.5), spanning log-spaced learning rates and regularization strengths together with discrete architectural choices (kernel width, embedding and attention dimensions, network size, and the number of sequences sampled per repertoire). Configuration C1 is each method’s benchmark default. Within each configuration, the per-method training budget is held fixed (up to 100 epochs with early stopping for ABMIL; DeepTCR’s own convergence criterion; 10,000 gradient updates for DeepRC).

Tuning proceeds in two stages. For Rawat T1D and CMV, all tuning uses folds 0 and 1 and the evaluation fold is never seen; for the seven 3-fold settings, configuration selection is not nested within the outer cross-validation (see below). *Stage 1* trains every configuration at seed 7 and ranks configurations by mean validation AUROC across the tuning contexts (Table B.6). *Stage 2* re-trains the top-ranked configuration at seeds 21 and 42 on the same contexts to measure its seed variability. The selected configuration is then retrained at all three seeds and evaluated with the benchmark protocol (Table B.7).

For the seven 3-fold settings (COVID-19, HIV, Influenza, Lupus, pooled T1D, RA, and TB), the tuning contexts rotate over the three benchmark folds: each trains on one fold, validates on a second, and leaves the third out. The selected configuration is then evaluated by standard 3-fold cross-validation, retrained on two folds and tested on the third. Because every fold serves as a validation fold in one tuning context, selection is not nested within the outer cross-validation, and the tuned estimates for these settings may be slightly optimistic. For Rawat T1D and CMV, tuning uses folds 0 and 1 only, and the selected configuration is evaluated once on the untouched fold 2 (the Rawat evaluation fold and the Emerson validation cohort), which is never used for selection.

### C.7 Encoder ablation protocol

To test whether pretrained sequence representations improve a deep MIL model, we replace the per-fold-trained CNN CDR3 encoder of ABMIL with one of three frozen pretrained encoders (Table B.8). ESM-2 embeddings are mean-pooled over last-layer residues, excluding padding and special tokens; TCR-BERT embeddings are mean-pooled from layer 8; TCR-VALID uses its 16-dimensional latent mean. ABMIL aggregation and the training protocol are unchanged. Results for all settings are in Table B.9.

### C.8 Sequencing-depth scaling protocol

For the sequencing-depth scaling analysis, we downsample each repertoire in both the training and test sets to a common target depth *D ∈ {*1k, 5k, 10k, 25k, 50k, 75k*}* unique sequences. To isolate the effect of depth from resampling noise, downsampling indices are pre-generated once per repertoire with a fixed seed and nested across depths: the *D* sequences used at depth *D* are always a subset of those used at any larger depth. We generate five independent downsampling replicates per depth and report mean AUROC (Figure 3a) and AUPRC (Figure A.2) with standard error across replicates. Methods that impose a default maximum sequencing-depth filter were overridden to accept the target depth *D*, so that all methods are evaluated on the same downsampled repertoires at each depth; Mal-ID is evaluated from 5k sequences upward.

The experiment is run on Lupus and HIV, chosen as representative diseases with distinct classification difficulty in the Mal-ID benchmark, and on five methods spanning distinct modeling paradigms: Emerson et al., DeepRC, VJ-kmer (Reg), VJ-kmer (XG), and Mal-ID. Specimens with fewer than 75k sequences are excluded so that all methods are compared on an identical specimen set across depths.

### C.9 Computational resource benchmarking

Computational resource usage was measured by running each method on a single NVIDIA L40S GPU (48 GB VRAM) with 50 CPU cores allocated per run. Computational benchmarking was run on one fold of the HIV dataset, encompassing data loading, featurization, model training, and evaluation. Wall-clock time was recorded using Python’s time.perf_counter() from process launch to termination. Peak CPU memory was estimated as the maximum resident set size (RSS), sampled from /proc/<pid>/status across the full process tree (including child processes) at 0.5-second intervals throughout execution. For GPU-accelerated methods, peak GPU memory was similarly polled at 0.5-second intervals via nvidia-smi, and a memory delta was computed by subtracting the pre-run GPU baseline to isolate method-specific allocation from ambient device usage. Each method was launched in an isolated process group, and a background sampling thread recorded peak values continuously until process exit. Total wall-clock time, peak RAM usage estimated as RSS, and peak GPU VRAM usage were recorded for each method.

### C.10 Ground Truth Driver Sequence Extraction

For each of HIV, COVID-19, and Influenza, we retained VDJdb entries [34] with confidence score *≥* 2, yielding 1,747 HIV, 2,355 COVID-19, and 36,453 Influenza reference CDR3s after deduplication within VDJdb. For COVID-19, we further augmented the reference set with 15,727 experimentally validated SARS-CoV-2-specific TCR*β* clonotypes from Minervina et al. [11], obtained using single-cell assays and restricted to specificity-confirmed binders. After deduplication against VDJdb, the final COVID-19 reference set contained 18,082 unique CDR3s.

### C.11 Driver sequence identification protocol

For models that support sequence-level ranking (either as prescribed by their authors or with minimal modifications), driver sequence identification was evaluated by using each trained disease classification model to assign a disease-association score to individual CDR3 sequences within held-out repertoires. For each repertoire, CDR3s were ranked by this score and compared against the groundtruth disease-associated CDR3 sequences present in each repertoire, as determined by the sequence extraction procedure described above. A repertoire CDR3 counts as a driver if it has at least 90% Levenshtein similarity to a reference CDR3 (Appendix C.12).

#### Emerson et al

We used the Fisher’s exact test statistics computed from training repertoires as sequence-level scores. Within each cross-validation fold, the p-value threshold was first selected on an internal train/validation split using validation AUROC, and TCR statistics were then recomputed on all non-test repertoires. Each test repertoire was represented as unique (V gene, J gene, CDR3 sequence) clonotypes, and each CDR3 was assigned the lowest Fisher’s exact test p-value observed among its matching (V, J, CDR3) tuples. CDR3s absent from the training-derived statistics were assigned a p-value of 1.0. CDR3s were ranked in ascending p-value order, and top-ranked sequences were compared against the ground-truth driver CDR3s for that repertoire.

#### GIANA

Driver sequence scores were derived from the reference clusters produced by the GIANA training procedure. Within each fold, training sequences were clustered to form reference clusters, and test sequences were assigned to these clusters in GIANA’s query mode. For each reference cluster, we computed the fraction of reference members originating from disease repertoires, excluding clusters with more than 100 reference TCRs. Each assigned test CDR3 received the disease fraction of its assigned cluster, while unassigned sequences or sequences assigned to excluded clusters received score zero. If the same CDR3 appeared in multiple cluster assignments, we retained the maximum disease-fraction score before ranking CDR3s in descending order.

#### DeepRC

Driver sequence scores were obtained from the trained sequence embedding and attention modules. Each CDR3 in a held-out repertoire was encoded using the same amino acid alphabet, positional features, and raw clone-count scaling used by the DeepRC dataloader, and then passed through the trained CNN sequence embedding network and attention network. We used the resulting attention logits to rank sequences, implemented as a softmax over all valid sequences in the repertoire. Duplicate CDR3s were aggregated by retaining the maximum softmax-normalized attention weight, and CDR3s were then ranked in descending order by this attention-derived score.

#### ABMIL

Sequence-level driver scores were taken from the trained gated-attention pooling module. For each held-out repertoire, all sequences were loaded without subsampling and converted to the model’s amino acid, V-gene, and J-gene tensor inputs. The per-sequence encoder produced instance embeddings, which were passed through the gated-attention branches to obtain one pre-softmax attention score per sequence. These scores were softmax-normalized across all sequences in the repertoire. Duplicate CDR3s were aggregated by retaining the maximum attention weight. CDR3s were then ranked by descending attention weight.

#### VJ-kmer (Reg)

Each unique (V gene, J gene, CDR3 sequence) tuple in a held-out repertoire was scored directly using the fitted logistic regression submodels. The k-mer submodel represented each CDR3 by its gapped 4-mer counts and applied the trained vectorizer, scaler, and logistic-regression decision_function, yielding a signed margin score. The V/J submodel represented each tuple by single-count V- and J-gene features and likewise applied the fitted vectorizer, scaler, and decision_function. The per-tuple driver score was the learned weighted combination of the margins from the two submodels, *αs*_kmer_ + (1 *− α*)*s*_VJ_. Duplicate CDR3s were aggregated by retaining the maximum score, and CDR3s were ranked in descending order.

#### VJ-kmer (XG)

Each unique (V gene, J gene, CDR3 sequence) tuple in a held-out repertoire was scored directly using the fitted XGBoost submodels. The k-mer submodel represented each CDR3 by its gapped 4-mer counts and used the trained XGBoost booster to produce a disease probability under the binary:logistic objective. The V/J submodel represented each entry by single-count V- and J-gene features and likewise produced a disease probability. The per-entry driver score was the learned weighted combination of these probabilities, *αp*_kmer_ + (1 *− α*)*p*_VJ_. Duplicate CDR3s were aggregated by retaining the maximum score, and CDR3s were ranked in descending order.

### C.12 Sequence and specimen overlap

#### Pooled T1D

The Mal-ID and Mitchell T1D specimens do not overlap: across all cross-source specimen pairs, the maximum CDR3 amino-acid Jaccard index is 0.013, and no pair exceeds 0.10, so no individual is duplicated across sources. Sequence overlap between the two sources is at the level expected for public clones (Mal-ID–Mitchell CDR3 amino-acid Jaccard index 0.035). The apparent asymmetry, in which *∼*31% of Mal-ID CDR3s appear in Mitchell but only *∼*4% of Mitchell CDR3s appear in Mal-ID, reflects Mal-ID’s shallower sequencing depth (Figure A.1): its smaller, more public-skewed sequence set is recoverable in Mitchell’s deeper pool, rather than indicating shared samples. This cross-protocol structure is what the pooled setting is designed to expose, and we do not consider it leakage.

#### Driver sequences

By construction, a driver is a repertoire CDR3 with at least 90% Levenshtein similarity to a reference CDR3, so overlap between repertoires and references is the measured signal rather than a confound. Driver-matched sequences make up 0.0004–0.03% (exact match) and 0.02–1.4% (fuzzy match) of the sequences in each disease repertoire, while *∼*40–50% of reference CDR3s are recovered by fuzzy match in the matching disease repertoires. The reference sets are fixed, identical for all methods, and never used as training labels.

### C.13 Experiment setup for demographic confounding analysis

#### Demographic-matched cohort

For each of the four Mal-ID diseases analyzed in Figure 5b—HIV, Lupus, Influenza, and COVID-19—we take a subset of the healthy repertoires matched on the cohort’s dominant confounder. For HIV, we apply a *symmetric ancestry filter*: both the disease and healthy rows are restricted to specimens with self-reported African ancestry, the dominant ancestry group in the HIV cohort. For the three age-confounded diseases, we apply an *age-match-healthy* rule: the disease cohort is left unchanged and Healthy/Background is subsampled so that its 10-year-binned age histogram has the same shape as the disease cohort’s. The matched healthy size *N* is the largest number consistent with the disease age histogram and the per-bin healthy availability; bins occupied by the disease but not the healthy pool are excluded. Each random-baseline cohort uses the *same disease side* as its matched run but replaces the healthy side with a uniformly random draw of the same *N* from the full healthy pool (ignoring demographics), isolating the effect of demographic structure from the cohort-size effect alone. We run five random-baseline replicates and report their mean AUROC; the matched cohort is a single draw (for HIV, a fixed ancestry filter). Within each (disease, cohort) configuration, classification follows the protocol of Section 4.1, reusing the preassigned 3-fold cross-validation splits. This analysis covers eight methods: Emerson et al., Ostmeyer et al., DeepRC, DeepTCR, ABMIL, VJ-kmer (Reg), VJ-kmer (XG), and Mal-ID.

#### Demographic-aware classification

For Figure 5c we evaluate three representative methods— DeepTCR, ABMIL, and VJ (Reg)—on eight datasets: HIV, Lupus, COVID-19, Influenza, TB, RA, CMV, and Rawat T1D. These three methods were chosen because each yields a fixed-length repertoire-level feature vector to which demographic covariates can be appended: V/J-gene usage frequencies in VJ (Reg), and the bag-level embeddings of ABMIL and DeepTCR, the two deep MIL methods with the highest mean AUROC in the main benchmark (Table B.1). Each model uses the demographic covariates recorded for its cohort: age and sex, plus self-reported ancestry (race/ethnicity for CMV) where recorded; ancestry is not recorded for Influenza, Rawat T1D, TB, or RA (Figure 5a). Within each panel, specimens missing any of these covariates (or with empty ancestry, where ancestry is used) are dropped before any model is fit, so that the *TCR only*, *TCR + demographics*, and *Demographics only* bars are evaluated on an identical specimen set (Table B.11 gives the resulting *n*). Demographic covariates are encoded as a numeric vector [age (raw), sex *∈ {*0, 1*}* (binary), one-hot ancestry], with the ancestry vocabulary fixed from the training fold and applied unchanged to the test fold. The *TCR + demographics* variant of each method appends this vector to the method’s repertoire-derived features and trains a single L1-penalized logistic regression on the concatenated representation. The repertoire features are V/J gene relative-frequency counts (the V/J sub-model of the VJ-kmer ensemble) for VJ (Reg), and the bag-level embeddings produced by a per-fold-trained ABMIL or DeepTCR model for the two deep-learning methods. Both feature blocks are standardized per fold before concatenation, and the corresponding *TCR only* bar reuses the same pipeline but drops the demographic block. The *Demographics only* baseline fits an L2-penalized logistic regression on the demographic feature vector alone, on the same specimen set. Rawat T1D is evaluated on held-out fold 2 and CMV on the Emerson validation cohort; all other datasets use the predefined 3-fold CV splits with pooled out-of-fold predictions, consistent with Section 4.1. We report AUROC and AUPRC with 95% participant-level bootstrap confidence intervals.

## Notes

### Summary of Updates

The new version includes author contributions.

